# Restoration of urban salmon habitat has limited effects on a key ecosystem function

**DOI:** 10.1101/2025.04.11.648418

**Authors:** Amanda R. Wik, Chelsea J. Little

## Abstract

Pacific Salmon have the potential to act as umbrella species in western North America. Urban streams have warmer temperatures, higher nutrient loads, and flashier hydrographs from impervious surroundings (urban stream syndrome). These changes are likely to affect ecosystem functions that are key to biodiversity and ecosystem services. Here, we ask whether restoration to improve urban stream habitat for Pacific Salmon can naturalize organic matter decomposition rates. Using structural equation models, we investigated the relationship between physical habitat, surrounding impervious surface, restoration intensity, and cotton strip decomposition rates, for streams in Vancouver, British Columbia, Canada. We hypothesized that 1) decomposition rates would differ along a gradient of urbanized to more natural reaches, and 2) sites receiving more intense restoration would have decomposition rates more similar to less-urbanized sites, partly because restoration would mitigate the environmental changes associated with urban stream syndrome. Streams surrounded by more impervious surface tended to have faster decomposition rates (standardized path coefficient = 0.38, the strongest direct effect on decomposition out of all the variables we considered). Reaches with higher restoration showed altered physical habitat characteristics compared to less-restored urban reaches. However, restoration intensity did not have a significant effect on decomposition rates in our structural equation model. Human impacts on freshwater ecosystems occur through pathways at broad spatial scales, and our results suggest that local efforts focusing on physical habitat restoration for umbrella species may not be the most effective way to address changes to ecosystem functioning.

## Introduction

Urbanization around streams can influence ecosystem functioning through a myriad of pathways, from impacts on the aquatic community to physical ecosystem structure and connectivity with terrestrial ecosystems. The urban stream syndrome refers to a consistent pattern of higher water temperatures, higher concentrations of nutrients and contaminants, and flashier hydrographs caused by surrounding impervious surface and channel modifications (Walsh et al. 2005). The increased water temperatures and nutrient loads affect the rates of many ecosystem processes such as decomposition and primary production (Sudduth et al. 2011, Woodward et al. 2012, Follstad Shah et al. 2017). Meanwhile, urbanization also alters stream morphology through channelization and diversion (Paul and Meyer 2001, Elmore and Kaushal 2008), reduction of large woody debris (Finkenbine et al. 2000), and decreased channel complexity (Paul and Meyer 2001, Walsh et al. 2005). These physical and chemical changes shift the total and relative abundance of species, functional groups, and key functional traits that contribute to ecosystem functions (Collier and Clements 2011, Marques et al. 2019, Classen-Rodríguez et al. 2019). Ecological restoration of urban streams aims to apply interventions such as daylighting, natural channel design, barrier removal, and habitat restoration to return freshwater populations, communities, or processes back toward a less-impacted state.

As highly valued species that are the target of many restoration activities in the Pacific Northwest, Pacific Salmon have the capability to act as “umbrella species” for stream ecosystems. The umbrella species concept postulates that conservation targeted at the habitat of one species or group of species provides benefits to co-occurring species (Kalinkat et al. 2017). Umbrella species are often charismatic or culturally important, and their ecological characteristics provide benefits to a wider ecosystem: extensive range and large body size, resulting in large areas being conserved; sensitivity to disturbance, requiring that conserved areas do not become degraded; and overlapping habitat requirements with co-occurring species. The life history and traits of Pacific Salmon make them excellent candidates for umbrella species because they are some of the largest freshwater predators in their ecosystems, they migrate over large distances, they co-occur with macroinvertebrates and other freshwater organisms, and they are sensitive to human disturbance (Branton and Richardson 2014, Warkentin et al. 2019). Although potential freshwater umbrella species generally receive less attention than terrestrial species (Branton and Richardson 2011, Kalinkat et al. 2017), restoration projects targeting their habitat could mitigate environmental degradation in human-impacted streams (Barnas et al. 2015, Walsh et al. 2020a).

Because the definition of umbrella species focuses on conservation of biodiversity, past assessments of potential freshwater umbrella species have focused on taxonomic richness (Bifolchi and Lodé 2005, Branton and Richardson 2011, 2014, Gilby et al. 2017). This strategy neglects the functional consequences of community composition and abundance: diverse communities often have higher rates of ecosystem functions, while higher-functioning ecosystems also typically support more biodiverse assemblages (Grace et al. 2016, Heilpern et al. 2018). We thus aimed to examine how habitat restoration targeting umbrella species in degraded urban ecosystems may affect ecosystem function, which is increasingly considered an important metric in ecosystem health assessments because it integrates biotic and abiotic pathways (Gessner and Chauvet 2002, Ranta et al. 2021).

Decomposition is a key ecosystem function that links terrestrial and aquatic portions of watersheds. Loss of detritivore diversity has been found to decrease rates of decomposition (Gessner et al. 2010). Considering that the urban stream syndrome is associated with lower invertebrate diversity, decomposition could be an effective tool for monitoring urban streams. Some urban streams have demonstrated increased rates of decomposition (Chadwick et al. 2006, Yule et al. 2015), while others have shown decreased rates (Chadwick et al. 2006, Rossi et al. 2019, Classen-Rodríguez et al. 2019), either of which can modify the flow of energy in aquatic food webs. Meanwhile, restoration efforts have been found to increase ecosystem functions, including leaf litter decomposition (Lepori et al. 2005, Frainer et al. 2018). When monitoring restoration outcomes, measuring decomposition has the added benefit of standardized assays that are relatively easy to implement and do not require long hours of identifying specimens like other biodiversity assessments (Tiegs et al. 2013).

To investigate the potential of Pacific Salmon as umbrella species, we tested whether restoration efforts aimed at conserving these species have benefits to the wider ecosystem, through organic matter decomposition rates measured using a standardized cotton strip assay. We measured the intensity of restoration interventions (Chazdon et al. 2021) rather than the abundance of the umbrella species, as regardless of the effectiveness of restoration for salmon returns, restoration could still provide benefits to the wider ecosystem. We also measured restoration intensity at both the local scale, directly surrounding sites where decomposition was measured, as well as across the watershed, reflecting that environmental conditions at broader spatial scales contribute to determining ecosystem processes measured locally. We predicted that in line with the urban stream syndrome, more urbanized sites would have higher water temperatures, higher nutrient concentrations, and altered physical habitat. We then hypothesized that 1) decomposition rates would be either higher (due to temperature, nutrients, and possibly habitat) or lower (due to contaminants or habitat alteration) in more urbanized reaches compared to more natural reaches, and that 2) sites where restoration interventions were more intense would have decomposition rates more similar to less-urbanized sites, because restoration would mitigate environmental degradation.

## METHODS

### Study area and sites

Vancouver is located on the west coast of British Columbia, Canada, and has a population of over 2.5 million, which has led to many of the impacts associated with the urban stream syndrome. Summer base flows and water velocity are both lower in Vancouver streams as compared to less urban counterparts, and impervious surface has caused higher turbidity and nitrate loads, along with lower dissolved oxygen (Finkenbine et al. 2000, Ruan et al. 2019). Vancouver’s streams historically provided passage to spawning anadromous salmonids, including Chinook, Coho, Sockeye, Pink, Chum, and Cutthroat, but in many streams, salmon populations dwindled and disappeared with increasing development pressures (Chen et al. 2017, Finn et al. 2021). In the last several decades, the cultural and economic values of salmon have led to a surge in efforts by governments, and most notably citizen Streamkeepers, to restore the streams (Pacific Streamkeepers Federation 2022). These efforts have ranged in scale from low-cost, volunteer-based riparian plantings and cleanups, to higher-cost professional culvert upgrades, and to major re-engineering of streams such as daylighting and channel reconstruction. We selected nine streams on Vancouver’s North Shore (Brothers Creek, Cypress Creek, Eagle Creek, Gallant Creek, McDonald Creek, Mosquito Creek, Parkside Creek, Rodgers Creek, and Westmount Creek) and one in the City of Vancouver (Still Creek) (Table 1), representing a gradient of urbanization and restoration (Table S1). All the streams pass through developed areas, with dense housing in the lower reaches. The upper reaches of the North Shore streams pass through forested areas or low-density housing. Two to five sample sites were chosen for each stream based on total stream length, the location of existing temperature loggers, proximity to restoration projects, and site accessibility. All but one stream had at least one reach restored. The chosen sampling reaches represented a combination of restored and unrestored reaches within watersheds, enabling comparisons among non-urbanized, urbanized restored, and urbanized unrestored reaches (Table S1). A total of 30 sites were chosen for this experiment (Fig. S1), although one was later discarded due to loss of sampling equipment.

**Table 1.** Profile of study streams with presence of the potential umbrella species, degree of urbanization, and restoration intensity undertaken across the watershed. Mean catch per unit effort is presented for a subset of the study streams, sampled in July of 2021, as well as the number of salmon stocked in the streams by enhancement programs (Gemmell, 2023).

| Stream | Species of salmon | Mean catch per unit effort | Stocked (number) | Average impervious surface (%) | Restoration intensity |
| --- | --- | --- | --- | --- | --- |
| Brothers Creek (BRT) | Chinook, Chum, Coho, Cutthroat, Pink, Rainbow, Steelhead | 0.144 | No | 27.90 | 16 |
| Cypress Creek (CPS) | Chum, Coho, Cutthroat, Pink | 0.432 | Yes (6,147) | 24.63 | 3 |
| Eagle Creek (EAG) | Chum, Coho, Cutthroat | 0.537 | Yes (1,401) | 12.98 | 12 |
| Gallant Creek (GLT) | Cutthroat | Not assessed | No | 9.04 | 10 |
| McDonald Creek (MCD) | Chum, Coho, Cutthroat, Steelhead | 0.144 | No | 35.27 | 26 |
| Mosquito Creek (MSQ) | Coho, Chum, Pink | 0.110 | No | 26.25 | 60 |
| Parkside Creek (PRK) | Coho, Cutthroat | Not assessed | No | 10.91 | 15 |
| Rodgers Creek (RGR) | Chum, Coho, Cutthroat, Pink | 0.400 | Yes (2,970) | 19.81 | 15 |
| Still Creek (STL) | Chum | Not assessed | No | 62.67 | 29 |
| Westmount Creek (WTM) | None | Not assessed | No | 30.79 | 0 |

### Decomposition and temperature measurements

We prepared cotton strips following methods from Tiegs et al. (2013), using Fredrix Unprimed Cotton Duck Canvas 15113 (Tara, Lawrenceville, Georgia, USA). We created 8 cm long, 27 thread wide cotton strips with 3 mm of fringe on the long edges and fringe 4 to 5 threads wide on the short edges. Cotton strips were deployed in 2021 for an early-summer study period on May 22, 23, and 28, and for a mid-summer study period on July 28 and 29. We deployed four cotton strips per site in each study period. To secure the strips in the stream, we used 20 cm square cement blocks with an open center, weighing 10 kg (Basalite, Surrey, B.C., Canada). We attached two cotton strips to each block by first securing a large zip tie around the block, and then attaching two cotton strips using smaller zip ties. Some sites had existing temperature loggers, otherwise we attached a HOBO Pendant Temperature/Light Logger UA-002-08 (Onset Computer Corporation, Bourne, Massachusetts, USA) to one concrete block per site, again using a small zip tie. One block was placed in a pool and the other in a riffle, except where this was not representative of the structure of the stream, in which case both may have been placed in pools or riffles. Blocks were placed to ensure they would remain fully submerged under water. For two sites, blocks were later moved due to high flow events and input from stakeholders.

Cotton strips were retrieved at the end of the early-summer study period on June 24 and 25, and at the end of the mid-summer study period on August 24 and 25. When retrieving cotton strips, we gently brushed the strips with a paint brush to remove organisms and particles. We then lowered the strips into Falcon tubes containing 70% ethanol for 30 seconds to remove microbial communities, and wrapped each strip in aluminum foil. Strips were frozen until the end of the collecting period, when they were placed in a drying oven at 40°C for 90 hours. Eight control samples were not deployed in streams, but were soaked in ethanol for 30 seconds prior to drying.

We measured the tensile strength of the cotton strips on an Instron 5469 Universal Testing Machine (Norwood, Massachusetts, USA). After drying, cotton strips were prepared for tensile strength measurement by painting 2 cm at the ends of the cotton strips with Acrylic Mediums Gloss Gel 5708 (Liquitex, Piscataway, New Jersey, USA) to prevent them from slipping out of machine grips. The machine was equipped with a 2 kN load cell and 50 kN large clamps (Cat. # 2716-020). The rate of advancement was set to 2 cm/min. Decomposition rates were calculated as the percent tensile loss per degree day (TLDD) using Eq. 1 (Tiegs et al. 2013),

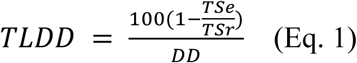

where TSe is the final tensile strength of the cotton strip and TSr is the average tensile strength of the control strips. Where this resulted in a negative number, TLDD was assigned to be zero.

Data from the temperature loggers was used to calculate degree days (DD): the average temperature for each day the loggers were deployed, summed over the deployment period. Three of the pre-existing temperature loggers malfunctioned during the deployment period: BRT02, MSQ01 and MSQ04. Because we found a high correlation between degree days and elevation, we predicted the degree days based on a linear relationship for those sites.

### Water chemistry

Water chemistry samples were taken during the early-summer study period on the same days as instrument deployment. The streams were sampled for pH, total ammonia and dissolved orthophosphate, and analysis was performed by ALS Environmental (Burnaby, B.C.). Water samples were kept in a cooler with ice packs and delivered to the lab by the end of the sampling day. Ammonia was measured by flow-injection analysis with fluorescence detection after reaction with orthophthaldialdehyde, and dissolved orthophosphate was measured colourimetrically after filtering through a 0.45 micron membrane filter. The samples were taken at 1 to 3 sites for each stream depending on its length, with one site near the outlet, the middle, and the headwaters to be representative of changes along the stream length.

### Habitat measurements

For the early-summer study period, we measured habitat and stream characteristics for each site in the three weeks following instrument deployment. The wetted width was measured across the existing channel flow, while the bankfull width was measured by estimating the furthest extent the channel could carry water at highest flow. Depth and flow velocity were measured using a FP111 Flow Probe (Global Water Resources, Phoenix, Arizona, USA) in representative pools and riffles. Using a 1 m^2^ quadrat divided into 25 squares, the substrate was characterized for a pool and a riffle into categories of impermeable surface, rocks (diameter > 20 cm), gravel (diameter 2.5-20 cm), fine gravel (diameter < 2.5 cm), sand, mud, living plants, leaf litter, biofilm, and woody debris. The percent pool and riffle cover were estimated for a stream reach encompassing the cement blocks (up to 5 meters long). We quantified canopy cover above the stream reach using a spherical crown densiometer, Convex Model A (Forest Densiometer, Marianna, Florida, USA). Finally, the riparian vegetation was designated into type (forest, field, meadow, housing) and dominant species of tree, shrub and ground cover (Table S2). A limited selection of habitat and stream characteristics that were predicted to change over the season were repeated for the mid-summer study period on the same day as instrument collection: wetted width, depth, and velocity.

### Impervious surface and restoration

To measure urbanization, we used Google Earth Pro to calculate a weighted average of the percent impervious surface (data source: Metro Vancouver 2019) of a 400 m^2^ quadrat around each site.

We collected information on restoration from news articles and Streamkeeper databases. We catalogued a total of 80 restoration projects that were undertaken between 1995 and 2020 in our study streams (Table S3). For our metric of restoration intensity, projects were assigned as either lightly-assisted (and given a score of 1 point), moderately-assisted (2 points), or heavily-assisted (3 points) in line with the suggested classification in Chazdon et al. (2021) (Table 2 and Table S3). The score per site was the sum of all the project scores that occurred within an approximately 400 m^2^ area surrounding each study site, while the score per watershed was the sum of the project scores for the entire stream. Restoration intensity per site ranged from 0 to 21, and per watershed from 0 to 62.

**Table 2.** Summary of Restoration Intensity metric, based on classifications from Chazdon et al. (2021)

| Restoration Intensity | Types of Restoration | Number of Projects in Category |
| --- | --- | --- |
| Lightly-assisted | Installing incubation boxes; cleaning-up in the stream and riparian areas; and removing invasive plants. | 6 |
| Moderately-assisted | Upgrading culverts (adding baffles and shortcrete, creating a natural bottom); adding diversion pipes to augment summer flow; adding fish ladders; adding timber guides; building box culverts; removing invasive plants and adding native riparian plantings; adding boulders and logs into the stream; installing debris fences and anchors; cleaning in-stream blockages. | 42 |
| Intensively-assisted | Restoring the stream mouth (adding series of pools and drops, redefining the estuary path); constructing rearing ponds; constructing or lengthening side channels; excavating plunge pools and adding boulders to raise the water level; bank stabilization and creek reconstruction to reduce sediment loads; removing channelization of the stream; daylighting the stream; widening the creek; reconstructing the streambed and adding habitat complexity; converting or restoring the riparian habitat to wetlands and meadows; installing bioswales; construction of weirs and rebuilding intakes; removing gravel from the stream. | 32 |

We developed a parallel metric of restoration effort to test the robustness of our results. To calculate restoration effort, each restoration project was assigned 1 point if it had a component within the stream and 1 point for a component in the riparian zone, and points were multiplied by the length of the project (in years). Restoration effort per site ranged from 0 to 15, and per watershed from 0 to 33. The two metrics were highly correlated (r_s_ = 0.91), but provided different information (Table S3). Because restoration effort is primarily a vote-counting method while restoration intensity incorporates qualitative information about restoration activities, our results here focus primarily on restoration intensity. However, we discuss differences between restoration intensity and restoration effort and show results based on restoration effort in the Supplemental Material.

### Statistical analysis

All statistical analyses were completed in R statistical environment version 4.4.1 (R Core Team 2024). Data was cleaned using packages ‘tidyr’ version 1.3.1 (Wickham 2020) and ‘dplyr’ version 1.1.4 (Wickham et al. 2020).

We examined the correlation between habitat variables using Spearman’s rank-order correlation. To assess the correlation between variables measured separately in early and late summer (e.g. stream depth, width, and velocity), we chose a positive association as the alternative hypothesis; for other correlation tests we used a two-sided test. To address our hypotheses, we needed to account for the fact that lower-elevation sites are much more urbanized (Spearman’s *ρ* = −0.51, p = 0.004), and that non-urbanized sites have not undergone restoration. To account for these correlations and assess direct and indirect effects of our factors of interest, we used a combination of linear mixed-effects models using the ‘lme4’ package version 1.1-35.5 (Bates et al. 2015) and piecewise structural equation models using the ‘piecewiseSEM’ package version 2.3.0.1 (Lefcheck 2016).

To assess whether streams surrounded by higher amounts of impervious surface exhibited key characteristics of urban stream syndrome we first sought to understand the relationship between elevation and stream temperature by fitting a linear mixed-effect model with average temperature over the deployment period as the response variable, elevation as a fixed factor, and sampling period and watershed as random factors. To address the effects of urbanization and restoration on stream temperature, we then used the residuals from the first model as the response variable in a linear model with impervious surface and restoration intensity as interacting fixed factors. We repeated this procedure for total ammonia, dissolved orthophosphate, and pH (but because these factors were only measured once, we omitted sampling period). For each response variable, if the mixed-effect model with elevation did not show a significant effect of elevation, then rather than using the residuals we expanded the mixed-effect model to include the main effect of elevation plus an interaction between impervious surface and restoration intensity. For mixed-effects models, we used the MuMIn package version 1.48.4 (Bartoń 2020) to calculate marginal (fixed-effects only) and conditional (entire model) R2 values, and the lmerTest package version 3.1-3 (Kuznetsova et al. 2017) to calculate F statistics and p-values associated with fixed effects.

After selecting variables that were not highly correlated, we performed Principal Component Analyses (PCAs) to explore multivariate stream habitat characteristics using the ‘stats’ package (R Core Team 2024). Finally, we examined potential indirect effects of urbanization and restoration on TLDD using structural equation models (SEMs). This allowed us to control for the pervasive effects of elevation, which is both associated with water temperature and land use patterns, and to assess how any perceived effects of restoration could be related to urbanization, given that only developed sites were restored. We chose the piecewise SEM approach because it accommodates random effects, which were important since we had multiple sites per watershed and multiple decomposition measurements per site. Thus, our structural equation model consisted of four linear mixed-effects models. We scaled and centered our data, then modeled TLDD as a function of surrounding impervious surface, restoration intensity, elevation, the first principal component of the habitat characteristics PCA, and pool/riffle placement, with site as a random factor. The first principal component axis was modeled as a function of elevation and restoration intensity, with watershed as a random factor. Impervious surface was modeled as a function of elevation, and restoration intensity was modeled as a function of both elevation and impervious surface, both with watershed as random factors, to account for the fact that non-urbanized sites are not restored. We checked whether the assumptions of each component model were met using the ‘DHARMa’ package version 0.4.6 (Hartig 2022); see Supplemental Methods for additional details on our model formulation and selection procedure. SEM fit was further assessed by considering Fisher’s C, and the marginal and conditional R2 values of each component model in the SEM.

To explore how choices about the method for measuring restoration affects our conclusions, we constructed three additional SEMs. The first used site-specific restoration effort instead of restoration intensity. The second used a modified subset of restoration effort, where we considered only in-stream habitat restoration and excluded projects that focused on fish passage (such as fish ladders) and the construction of spawning habitat outside of the main stream channel (such as ponds).

Finally, in our main model we used TLDD as our metric of decomposition, because this allows a standardized comparison across sites with different water temperatures, and because temperature-adjusted decomposition rates are commonly found in other studies. However, to more explicitly examine the effect of temperature on decomposition rates, we built a fourth SEM which used the percent tensile loss per day (TLND; not temperature-adjusted) as the response variable of interest. Here, we included a component model with degree days as a function of elevation and impervious surface, with watershed as a random factor, and we adjusted the model of TLND to also include degree days.

## RESULTS

### Stream characteristics

The majority of the streams had a cascade morphology and were forested, with alder (*Alnus rubra*) and Western redcedar (*Thuja plicata*) as the dominant tree species (both native to the region). Streams had an average wetted width of 5 meters (range: 0.7-9.5 m), depth of 23 centimeters (range: 7-54 cm), and velocity of 0.34 meters per second (range: 0.1-3.6 m/s) in early summer. Variables that were measured separately for each sampling period tended to be very highly correlated, for example depth (r_s_ = 0.55, p = 0.002), wetted width (r_s_ = 0.86, p < 0.001), and temperature accumulated over the sampling period (r_s_ = 0.50, p = 0.003), with the notable exception of stream velocity (r_s_ = −0.16, p = 0.80).

### Temperature and nutrients

Average water temperature during deployments had a strong negative relationship with elevation (linear mixed-effects model, F_1,50.18_ = 27.47, p < 0.001; Fig. 1), and no further association with impervious surface, restoration or their interaction (when considering restoration intensity: F_3,54_ = 0.023, p > 0.7; when considering restoration effort: F_3,54_ = 1.28, p > 0.25). We confirmed that the effect of elevation on temperature was not biased by low-elevation streams being more urbanized (which is shown in Fig. S2), by repeating the linear mixed-effects model only with the portion of sites which were surrounded by ∼20% impervious surface or less (which spanned an elevational gradient of 14–394 meters above sea level), obtaining nearly identical results (linear mixed-effects model, F_1,13.43_ = 13.17, p = 0.003; estimated coefficient for the effect of elevation = −0.009 for both full and reduced datasets). Neither total ammonia (F_1,12.72_ = 0.01, p > 0.9; marginal R^2^ < 0.001) nor dissolved orthophosphate (F_1,12.63_ = 0.05, p > 0.8; marginal R^2^ = 0.001) showed associations with elevation, thus we expanded the linear mixed-effects models to include impervious surface and restoration intensity. For ammonia, the resulting model explained a large proportion of variance (conditional R^2^ = 0.43) but most of the variance was associated with the watershed random effect, rather than the fixed effects (marginal R^2^ = 0.05; all p-values for interactive and main effects > 0.05). Results were nearly identical for orthophosphate. By contrast, pH was strongly negatively associated with elevation (F_1,9.86_ = 93.04, p < 0.001; Fig. 1). In the linear model with the residuals as a response variable, there were no associations with impervious surface, restoration intensity, or their interaction (F_3,17_ = 0.87, p > 0.4).

**Figure 1.**
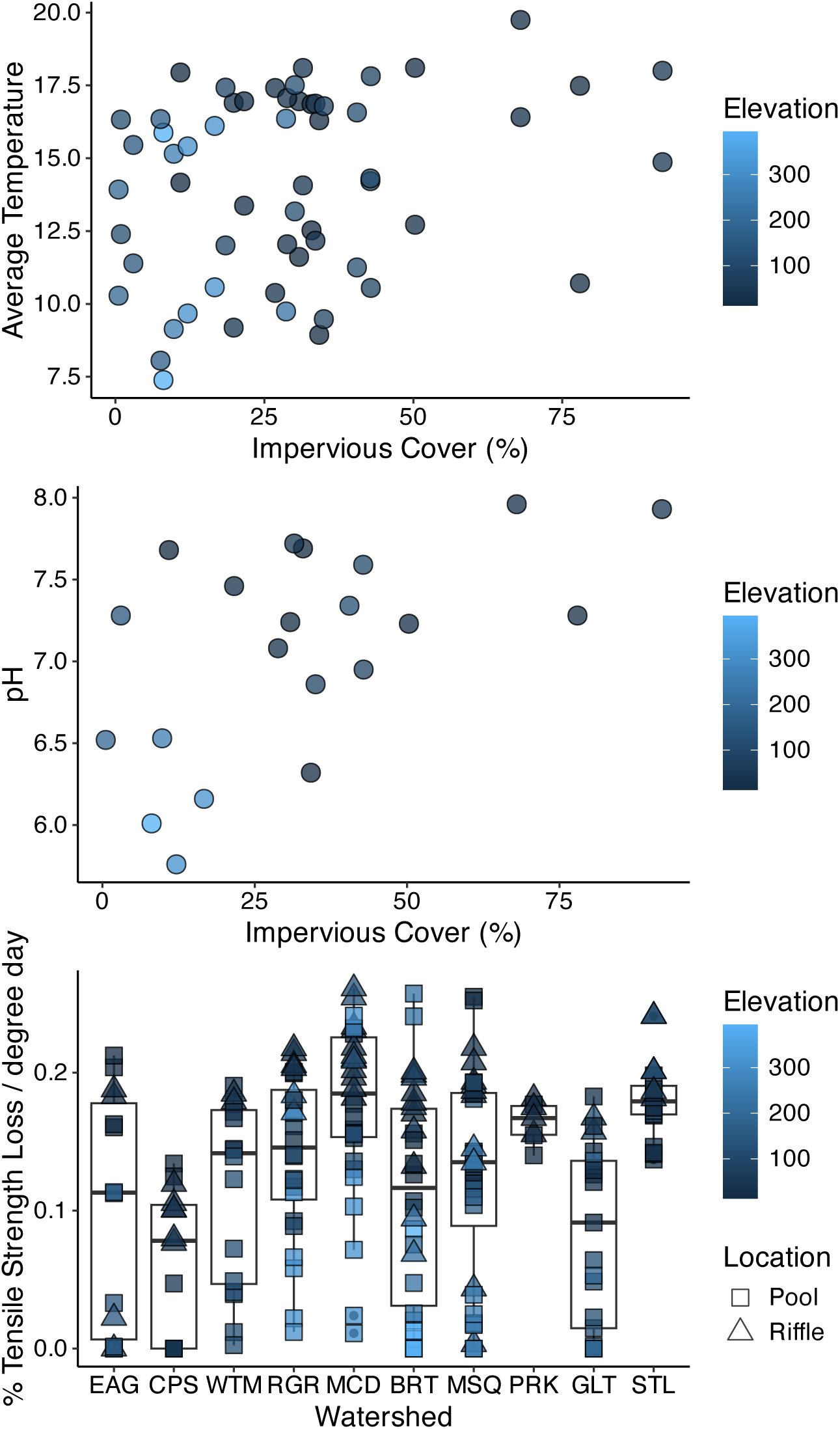
Key physical conditions, as well as decomposition rates, varied with elevation within and among watersheds. Average water temperatures during cotton strip deployment periods (a) and early-summer pH (b) both tended to be higher at sites surrounded by more impervious surface, which themselves were concentrated at low elevations. (c) Decomposition rate, measured as the percent of a cotton strip’s tensile strength which was lost per degree day during the deployment period, varied among streams and often with elevation within streams. In panel (c), streams are presented from west to east in the study area.

### Multivariate characterization of stream habitat

We found high correlations between several pairs of habitat variables, particularly those within the categories of water chemistry, microhabitat and temperature, which informed our decisions about which habitat variables to include in the PCA. We used only the June measurements of depth and wetted width; we excluded bankfull width completely because it was highly correlated with wetted width (from early summer measurements, r_s_ = 0.93, p < 0.001). The PCA explained 43% of the variation in the dataset with the first two principal component axes (Fig. 2). The first principal component explained 22.4% of the variance in the dataset, with strong positive loadings from percent pool, streambed sand and biofilm, while streambed rocks, June stream velocity, and overstory density had strong negative loadings. The second principal component explained 20.7% of the variance in habitat with strong positive loadings from wetted width, June stream depth, June stream velocity, and rocks, and strong negative loadings from fine gravel and mud streambed cover.

**Figure 2.**
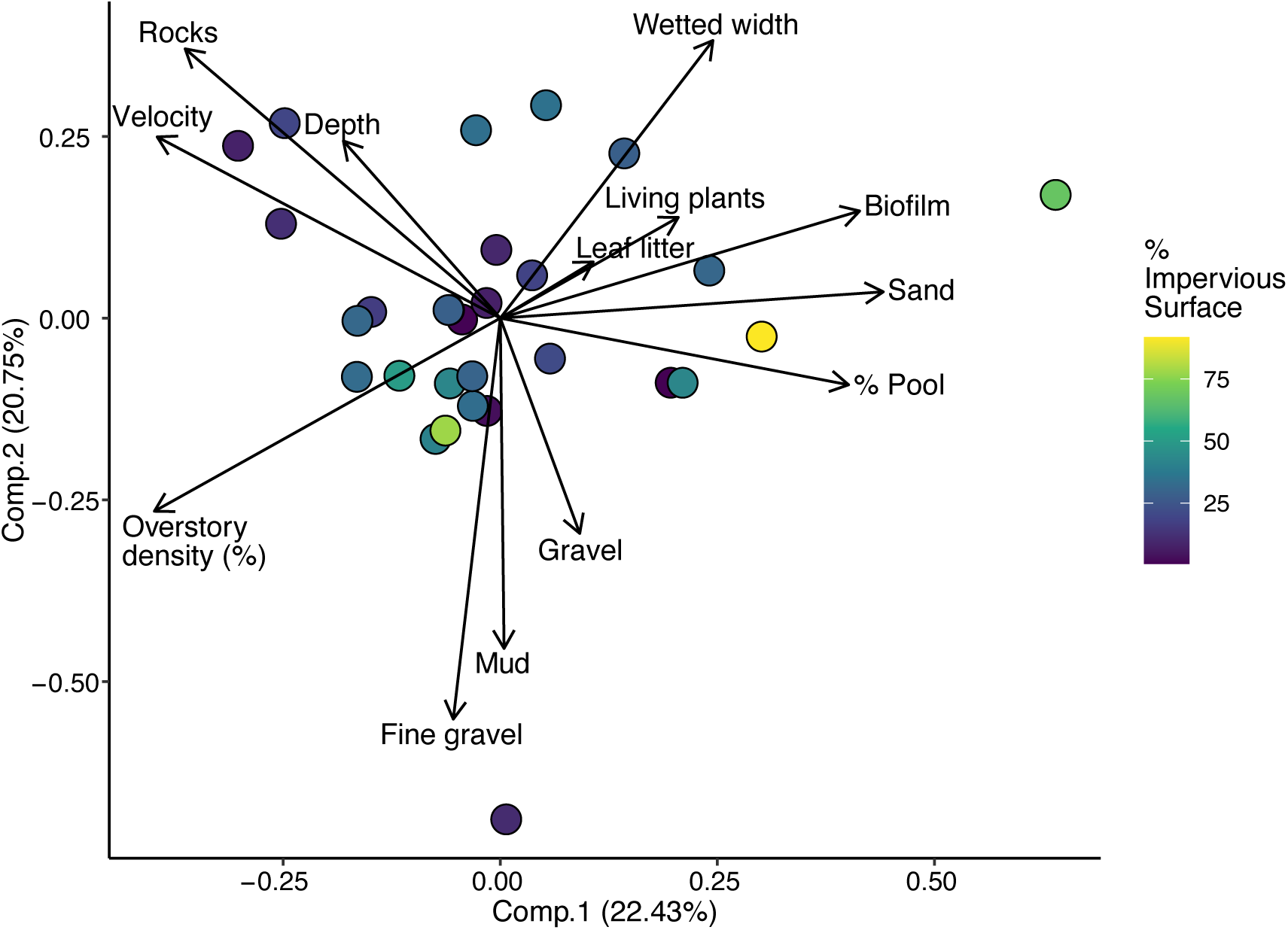
Principal component analysis of habitat variables measured at 28 study sites with complete data from the early summer study period.

A PCA of habitat characteristics at 21 sites where water chemistry measurements were collected was very similar (explaining 47.2% of the variation in the smaller dataset), and pH had a positive loading on the first principal component and a negative loading on the second principal component (Fig. S3).

### Overall effects of urbanization and restoration on decomposition rates

Decomposition rates (i.e. loss of tensile strength, TLDD) of 212 cotton strips collected from our study sites across the study periods varied from 0 to 2.97% per degree day, with a mean of 1.59% (Fig. 1). Piecewise SEM fit was only moderate (Fisher’s C = 17.804, p = 0.02; see Supplemental Material for discussion of model fitting); our SEM structure was nearly saturated, complicating the calculation of ÷^2^ and C, however we did not remove model components to increase directed separation because all paths were relevant to our specific hypotheses. Conditional R^2^ values were high (> 0.60) for each of the four component models, and marginal R^2^ (attributed to fixed effects only) ranged from 0.13 for the model of PCA principal component 1 to 0.43 for the model of TLDD.

In our piecewise SEM (Fig. 3), impervious surface had a direct, positive effect on decomposition (standardized coefficient = 0.38, p = 0.004), whereby cotton strips decomposed more quickly in urbanized sites with more impervious surroundings. Contrary to our hypotheses, restoration intensity did not have a strong effect on cotton strip decomposition rates (standardized coefficient [direct effect] = 0.15, p = 0.19). The main mechanism through which stream habitat influenced decomposition rate was via faster decomposition in pool versus riffle location (standardized coefficient [direct effect] = 0.25, p < 0.001). The first principal component of the habitat PCA, representing a gradient from canopy-covered, rocky reaches with fast-moving water, to sandy, biofilm-covered reaches, had no additional effect (standardized coefficient = −0.07, p > 0.5). As expected, elevation was important in our model of decomposition, but this was as an indirect rather than direct effect (Fig. 3). Elevation was negatively associated with impervious surface (standardized coefficient [direct effect] = −0.51, p < 0.001) and restoration intensity (standardized coefficient [direct effect] = −0.51, p = 0.001). Thus, elevation had an indirect effect on decomposition rate via impervious surface of −0.19, calculated from the component direct effects, whereby cotton strips deployed at lower elevations tended to decompose more quickly due to higher urbanization (Fig. 3).

**Figure 3.**
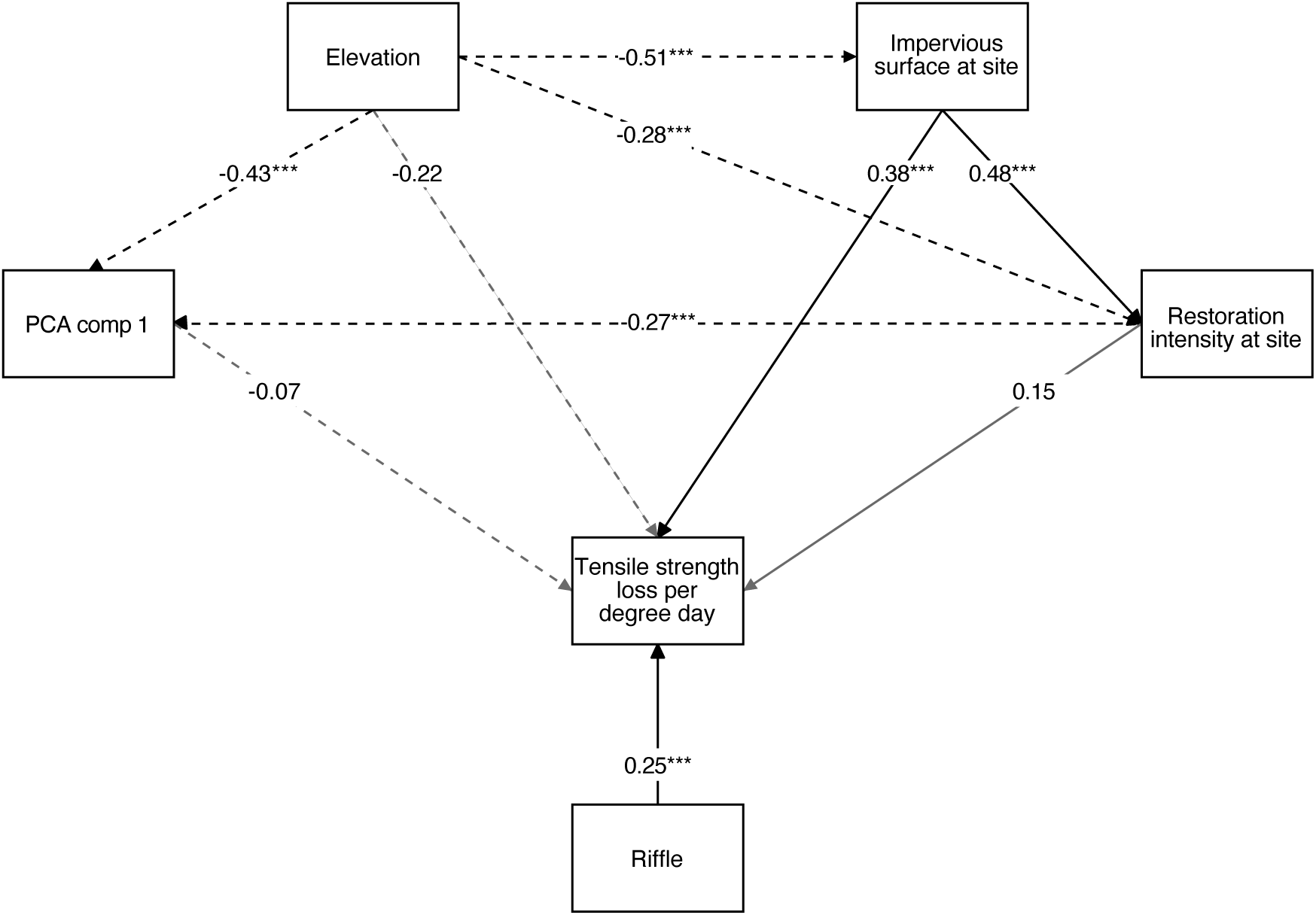
Results of a piecewise structural equation model (SEM) examining the direct and indirect effects of elevation, urbanization (as measured by amount of impervious surface surrounding a stream reach), restoration intensity at each stream reach, and habitat characteristics on cotton strip decomposition rates (measured as tensile strength loss per degree day) across the early- and mid-summer study periods. Significant (p < 0.05) paths are shown with black lines, and non-significant paths (p > 0.05) are shown with grey lines. Negative paths are indicated by dashed lines. Numbers in each path show the standardized path coefficients from the SEM. Results of a similar SEM on tensile strength loss per day, with temperature as a variable in the model, were qualitatively similar (see Fig. S4) and showed a direct and indirect (via temperature) influence of elevation on decomposition.

Piecewise SEM models using restoration effort rather than restoration intensity did not have improved global goodness-of-fit. Using complete restoration effort (Fisher’s C = 25.49, p = 0.001; Fig. S4), the direct effect of elevation became more important for TLDD, with cotton strips at lower elevations decomposing faster (standardized coefficient [direct effect] = −0.23, p = 0.05). Using restoration effort with only stream habitat restoration projects (Fisher’s C = 41.15, p < 0.001; Fig. S5) revealed a stronger effect of impervious surface on restoration effort (standardized coefficient [direct effect] = −0.26, p = 0.003) than the prior two models, but still no effect of restoration effort on TLDD (standardized coefficient [direct effect] = 0.11, p = 0.30).

Finally, building an alternative piecewise SEM focused on TLND and incorporating direct and indirect effects of temperature demonstrated somewhat improved global goodness-of-fit (Fisher’s C = 19.81, p > 0.10, but issues remained with calculation of ÷^2^). As expected, temperature had a strong direct effect on daily decomposition rates, and was colder at high elevations and in areas with less impervious surface, adding indirect effects of those variables on decomposition (Fig. S5).

## DISCUSSION

We measured organic matter decomposition in urban streams to investigate whether Pacific Salmon serve as umbrella species by providing the motivation for stream restoration projects that improve ecosystem function. Supporting our initial hypothesis, we found that organic matter decomposed faster in streams surrounded by more impervious surface, although it was difficult to disentangle the causal effect of urbanization from an underlying altitudinal gradient. We had also predicted that urbanized sites with more intense restoration would have similar decomposition rates to less-urbanized sites, however, we found little support for this hypothesis. Overall, we did not find strong evidence that restoration of Pacific Salmon habitat pushed decomposition rates back towards their presumed baseline.

There are multiple pathways through which urbanization is likely to affect decomposition. Here, we used the amount of impervious surface around a stream as a metric of urbanization. The influence of impervious surface on organic matter decomposition varies within the literature (Chadwick et al. 2006, Imberger et al. 2010, Kielstra et al. 2019). We found a strong, positive relationship whereby even after adjusting for the warmer water temperatures in streams surrounded by higher amounts of impervious surface, these streams had higher decomposition rates per growing degree day. This pattern could be explained by other elements of the urban stream syndrome, such as higher nutrient loads, which typically accelerate decomposition (Woodward et al. 2012). However, after controlling for the effect of elevation, we did not find evidence of higher concentrations of ammonia and orthophosphate in more-compared to less-urbanized stream reaches. Another possibility is that more urbanized reaches have different biological communities, and that this contributes to the additional effect on decomposition rates outside of the temperature pathway. Unfortunately, surveying biological communities was outside the scope of this project.

In addition to the number and type of restoration interventions, our study streams varied in their salmon stocking status (Table 1), which affects demography and expected adult returns. The effectiveness of umbrella species conservation has most commonly been assessed by comparing overall species richness in areas with varying abundances of the umbrella species (Roberge and Angelstam 2004, Branton and Richardson 2011). However, due to stocking in our study streams, disentangling the effects of restoration on salmon status, and then salmon status on communities and ecosystems, would be particularly complex. Thus, rather than assessing the abundance of the umbrella species, we considered how effort expended to protect them could benefit ecosystems, regardless of whether the desired effect on the focal species is achieved. Our approach represents a more expansive mode of considering “umbrella species” (i.e., Pacific Salmon) in two ways. First, we included the potential functional benefits to ecosystems rather than focusing solely on biodiversity. Many ecosystem functions are promoted by higher levels of biodiversity, therefore we expected that conservation of umbrella species would protect ecosystem functions. Second, we chose to more mechanistically examine the potential pathway from umbrella species to ecosystem function by focusing on habitat modification targeted at improving conditions in degraded ecosystems, rather than habitat protection, which assumes that ecosystems are intact and can be maintained that way.

Contrary to our hypotheses, we did not find evidence that restoration of salmon habitat resulted in different ecosystem functioning when compared to sites where less intense restoration efforts were undertaken. This could be for at least three reasons. First, it is possible that restoration did not change the stream environment. Our SEM did show a significant relationship between restoration intensity and the first principal component of habitat, which was characterized by wide, sandy reaches with prevalent pools, leaf litter and biofilm; this relationship was negative, suggesting that restored reaches were narrower, deeper, rockier, and with faster-moving water. Our SEM also showed that riffles had faster decomposition rates than pools, a finding consistent with previous decomposition studies, including those using cotton strips (Webb et al. 2019, Banks et al. 2023). Thus, this first possible explanation seems inconsistent with our data, although anthropogenic factors influencing stream structure and function often operate across broader scales than can be effectively addressed with reach-scale restoration projects (Bernhardt and Palmer 2011) and may have overwhelmed the local changes in specific physical conditions.

A second possible scenario for the lack of restoration effects on decomposition is that salmon may not be an umbrella species for stream biodiversity, and thus their conservation only minimally impacts biotically-mediated ecosystem function. Previous research reported that protecting habitat for predicted umbrella species does not necessarily result in protecting the type of habitat needed by other species of concern (Branton and Richardson 2014, Wang et al. 2021). In our case, this could be due to the types and locations of restoration projects that were undertaken. In the site with the highest restoration intensity (the outlet of McDonald Creek), these efforts focused mainly on the physical structure of salmon habitat and connectivity, including installing fish ladders and installing baffles around culverts. These types of projects are important for increasing salmon populations by reducing barriers to migratory pathways, but do not necessarily address habitat conditions that contribute to diversity or abundance of other stream biota. The main mechanism through which these lower-intensity, migration-focused projects may serve as umbrella interventions for other taxonomic groups is through eventual ecosystem-scale effects of the Pacific Salmon returns (Walsh et al. 2020b), should the projects have their intended benefits. On the other hand, projects that restore stream habitat rather than only restoring connectivity for fish movement have been shown to impact both the invertebrate assemblages responsible for organic matter processing and decomposition itself (Frainer et al. 2018, Muotka et al. 2002). Habitat restoration often falls under the intensively-assisted category, requiring expensive engineering solutions. Thus, while removal of the barriers that currently render many kilometers of stream length inaccessible to salmon (Finn et al. 2021) is a winning strategy from a cost-benefit perspective, salmon may be more effective umbrella species (e.g. over shorter timescales) for biodiversity and ecosystem processes when their protection motivates habitat restoration. We suggest that as barriers are removed and salmon can increasingly regain access to urban streams, more intensive habitat restoration approaches should have a role in ensuring habitat quality for not only salmon but also invertebrates and microbes that play key roles in stream ecosystem function.

Another reason that salmon may not be effective umbrella species in our study is due to the location of restoration projects within the stream network. Several sites with the highest restoration intensities were stream outlets. Restoration of physical structure and downstream reaches is an essential step in restoring salmon access to watersheds, but does not address the broader-scale anthropogenic impacts happening upstream and propagating downhill in urban watersheds, and thus may limit the positive effects on stream biodiversity. Elsewhere, research has shown that restoration efforts at the headwaters and in small watersheds have the highest impacts on ecosystem processes, because benefits from restoration efforts in downstream reaches may be overwhelmed by cascading effects from upstream (Levi & McIntyre 2020). On the other hand, Vancouver’s development pattern where steep slopes and provincial land ownership prevents urbanization at the top of watersheds presents a unique context for evaluating the scale of urbanization and restoration impacts. In our study area and in other cities with similar topography, headwaters are rarely urbanized, which justifies the restoration focus on downstream reaches.

Finally, a third possible scenario is that even if conservation of Pacific Salmon does protect other biodiversity, this would only promote ecosystem functioning if there is a strong, positive relationship between biodiversity and ecosystem function. In fact, the relationship between biodiversity and decomposition in freshwater ecosystems is not straightforward: abiotic and biotic conditions mediate the strength and slope of diversity-decomposition relationships (Lecerf and Richardson 2010, Beaumelle et al. 2020). For example, a recent study considering effects of land use on both freshwater biodiversity and cotton strip decomposition found strong effects on the former but not the latter (Koivunen et al. 2023), indicating a nonlinear relationship between biodiversity and decomposition.

Our conclusions about whether Pacific Salmon are effective umbrella species for stream ecosystems is likely related to the differing approach of our research compared to past work. We assessed the effectiveness of the umbrella species concept where a restoration scheme has actually been applied, as opposed to many studies that assess the diversity of co-occurring species across the umbrella species’ existing range without considering variation between conserved habitats and non-conserved habitats. As habitat loss continues globally, it will be important to understand whether the umbrella species concept works where habitat is already degraded, or if its primary utility is to prevent the conversion of intact habitat.

Yet, while Pacific Salmon may not have served as umbrella species for decomposition in our study, their presence in urban streams still has many benefits. As charismatic species, they help people connect with the natural world and generate interest in habitat protection and restoration, which is likely to have long-term benefits for ecosystems and their inhabitants far beyond the scale of a local restoration project. Future studies should incorporate biodiversity monitoring to understand whether the effects of restoration are mediated by biological communities that co-occur with the umbrella species. In a world where funding for environmental conservation and restoration is limited, there is an urgent need to prioritize actions that provide benefits to more than just one taxonomic group. Stream restoration is a key strategy for safeguarding Pacific Salmon (Walsh et al. 2020a), and could provide widespread benefits for biodiversity and ecosystem processes if we can identify which particular restoration interventions are effective at promoting ecosystem functioning.

## ACKNOWLEDGEMENTS

Author contributions: A.R.W. and C.J.L. conceived the ideas, designed the methodology and collected the data. A.R.W. performed initial analyses and wrote the first draft of the manuscript. C.J.L. performed additional statistical analyses. Both authors revised the manuscript and gave final approval for publication.

We acknowledge that our research was carried out on the traditional, unceded territories of the Coast Salish people including the Musqueam, Squamish, and Tsleil-Waututh Nations. We thank John Barker from the West Vancouver Streamkeepers, Andrew Egan from Langara College, and Adrian Avendaño from the Still Moon Arts Society for assistance with site selection and information about restoration; Nikki Kroetsch for sharing temperature data; Carmen Gemmell for sharing salmon data; and John Richardson for comments on an earlier version of this manuscript. We also thank Saeid Soltanian at the University of British Columbia Centre for Flexible Electronics and Textiles for training on the Materials Testing System. This project was funded by a Discovery Grant (to C.J.L.) and an Undergraduate Student Research Award (to A.R.W.) from the Natural Sciences and Engineering Council of Canada.

The authors declare no conflict of interest.

## Open science statement

Code used to perform analyses and generate figures is on GitHub at https://github.com/chelseajlittle/salmon-stream-decomp. Data will be made publicly available upon acceptance in a journal.

## Notes

### Competing Interest Statement

The authors have declared no competing interest.

## LITERATURE CITED

Banks, L. K., Lavoie, I., Robinson, C. E., and A. G. Yates. (2023) Effects of groundwater inputs on algal assemblages and cellulose decomposition differ based on habitat type in an agricultural stream. Hydrobiologia 850:3517–3537.

Barnas, K. A., S. L. Katz, D. E. Hamm, M. C. Diaz, and C. E. Jordan. 2015. Is habitat restoration targeting relevant ecological needs for endangered species? Using Pacific Salmon as a case study. Ecosphere 6:art110.

Bartoń, K. 2020. MuMIn: Multi-Model Inference. https://CRAN.R-project.org/package=MuMIn.

Bates, D., Mächler, M., Bolker, B., and S. Walker. 2015. Fitting linear mixed-effects models using lme4. Journal of Statistical Software 67:1–48.

Beaumelle, L., F. De Laender, and N. Eisenhauer. 2020. Biodiversity mediates the effects of stressors but not nutrients on litter decomposition. eLife 9.

Bernhardt, E. S., and M. A. Palmer. 2011. River restoration: the fuzzy logic of repairing reaches to reverse catchment scale degradation. Ecological Applications. 21:1926–1931.

Bifolchi, A., and T. Lodé. 2005. Efficiency of conservation shortcuts: an investigation with otters as umbrella species. Biological Conservation 126:523–527.

Branton, M. A., and J. S. Richardson. 2014. A test of the umbrella species approach in restored floodplain ponds. Journal of Applied Ecology 51:776–785.

Branton, M., and J. S. Richardson. 2011. Assessing the value of the umbrella-species concept for conservation planning with meta-analysis. Conservation Biology. 25:9–20.

Chadwick, M. A., D. R. Dobberfuhl, A. C. Benke, A. D. Huryn, K. Suberkropp, and J. E. Thiele. 2006. Urbanization affects stream ecosystem function by altering hydrology, chemistry, and biotic richness. Ecological Applications 16:1796–1807.

Chazdon, R.L., Falk, D.A., Banin, L.F., Wagner, M., Wilson, S.J., Grabowski, R.C., & Suding, K.N. (2021) The intervention continuum in restoration ecology: rethinking the active-passive dichotomy. Restoration Ecology e13535. DOI: 10.1111/rec.13535.

Chen, H., M. Chen, Q. Liu, and Y. Jiang. 2017. Vancouver Lost Stream Restoration Assessment. University of British Columbia. https://blogs.ubc.ca/jacksoneportfolio/files/2017/04/Vancouver-Lost-Stream-Restoration-Assessment.pdf.

Classen-Rodríguez, L., P. E. Gutiérrez-Fonseca, and A. Ramírez. 2019. Leaf litter decomposition and macroinvertebrate assemblages along an urban stream gradient in Puerto Rico. Biotropica 51:641–651.

Collier, K. J., and B. L. Clements. 2011. Influences of catchment and corridor imperviousness on urban stream macroinvertebrate communities at multiple spatial scales. Hydrobiologia 664:35–50.

Elmore, A. J., and S. S. Kaushal. 2008. Disappearing headwaters: patterns of stream burial due to urbanization. Frontiers in Ecology and the Environment 6:308–312.

Finkenbine, J. K., J. W. Atwater, and D. S. Mavinic. 2000. Stream health after urbanization. Journal of the American Water Resources Association 36:1149–1160.

Finn, R. J. R., L. Chalifour, S. E. Gergel, S. G. Hinch, D. C. Scott, and T. G. Martin. 2021. Quantifying lost and inaccessible habitat for Pacific salmon in Canada’s Lower Fraser River. Ecosphere 12.

Follstad Shah, J. J., J. S. Kominoski, M. Ardón, W. K. Dodds, M. O. Gessner, N. A. Griffiths, C. P. Hawkins, S. L. Johnson, A. Lecerf, C. J. LeRoy, D. W. P. Manning, A. D. Rosemond, R. L. Sinsabaugh, C. M. Swan, J. R. Webster, and L. H. Zeglin. 2017. Global synthesis of the temperature sensitivity of leaf litter breakdown in streams and rivers. Global Change Biology 23:3064–3075.

Frainer, A., L. E. Polvi, R. Jansson, and B. G. McKie. 2018. Enhanced ecosystem functioning following stream restoration: The roles of habitat heterogeneity and invertebrate species traits. Journal of Applied Ecology 55:377–385.

Gemmell, C. 2023. Interconnected effects of water temperature and habitat on relative abundance of juvenile salmonids during the summer months in coastal urban streams. M.R.M. Thesis, Simon Fraser University, Burnaby, BC.

Gessner, M. O., and E. Chauvet. 2002. A case for using litter breakdown to assess functional stream integrity. Ecology 12:498–510.

Gessner, M. O., C. M. Swan, C. K. Dang, B. G. McKie, R. D. Bardgett, D. H. Wall, and S. Hättenschwiler. 2010. Diversity meets decomposition. Trends in Ecology & Evolution 25:372–380.

Gilby, B. L., A. D. Olds, R. M. Connolly, N. A. Yabsley, P. S. Maxwell, I. R. Tibbetts, D. S. Schoeman, and T. A. Schlacher. 2017. Umbrellas can work under water: Using threatened species as indicator and management surrogates can improve coastal conservation. Estuarine, Coastal and Shelf Science 199:132–140.

Grace, J. B., T. M. Anderson, E. W. Seabloom, E. T. Borer, P. B. Adler, W. S. Harpole, Y. Hautier, H. Hillebrand, E. M. Lind, M. Pärtel, J. D. Bakker, Y. M. Buckley, M. J. Crawley, E. I. Damschen, K. F. Davies, P. A. Fay, J. Firn, D. S. Gruner, A. Hector, J. M. H. Knops, A. S. MacDougall, B. A. Melbourne, J. W. Morgan, J. L. Orrock, S. M. Prober, and M. D. Smith. 2016. Integrative modelling reveals mechanisms linking productivity and plant species richness. Nature 529:390–393.

Hartig, F. 2022. DHARMa: Residual Diagnostics for Hierarchical (Multi-Level / Mixed) Regression Models. https://CRAN.R-project.org/package=DHARMa

Heilpern, S. A., B. C. Weeks, and S. Naeem. 2018. Predicting ecosystem vulnerability to biodiversity loss from community composition. Ecology 99:1099–1107.

Imberger, S. J., R. M. Thompson, and M. R. Grace. 2010. Searching for effective indicators of ecosystem function in urban streams: assessing cellulose decomposition potential. Freshwater Biology 55:2089–2106.

Kalinkat, G., J. S. Cabral, W. Darwall, G. F. Ficetola, J. L. Fisher, D. P. Giling, M.-P. Gosselin, H.-P. Grossart, S. C. Jähnig, J. M. Jeschke, K. Knopf, S. Larsen, G. Onandia, M. Pätzig, W.-C. Saul, G. Singer, E. Sperfeld, and I. Jarić. 2017. Flagship umbrella species needed for the conservation of overlooked aquatic biodiversity. Conservation Biology 31:481– 485.

Kielstra, B. W., J. Chau, and J. S. Richardson. 2019. Measuring function and structure of urban headwater streams with citizen scientists. Ecosphere 10:e02720.

Koivunen, I., Muotka, T., Jokikokko, M., Virtanen, R., and J. Jyväsjärvi. 2023. Downstream impacts of peatland drainage on headwater stream biodiversity and ecosystem functioning. Forest Ecology and Management 543:121143.

Kuznetsova, A., P. B. Brockhoff, and R. H. B. Christensen. 2017. lmerTest Package: Tests in Linear Mixed Effects Models. Journal of Statistical Software 82:1–26.

Lecerf, A., and J. S. Richardson. 2010. Biodiversity-ecosystem function research: Insights gained from streams. River Research and Applications 26:45–54.

Lefcheck, J.S. 2016. piecewiseSEM: Piecewise structural equation modeling in R for ecology, evolution, and systematics. Methods in Ecology and Evolution 7:573–579.

Lepori, F., D. Palm, and B. Malmqvist. 2005. Effects of stream restoration on ecosystem functioning: detritus retentiveness and decomposition. Journal of Applied Ecology 42:228–238.

Levi, P. S., and P. B. McIntyre. 2020. Ecosystem responses to channel restoration decline with stream size in urban river networks. Ecological Applications 30:e02107.

Marques, P. S., L. R. Manna, R. Mazzoni, and R. El-Sabaawi. 2019. Intraspecific trait variation in urban stream ecosystems: toward understanding the mechanisms shaping urban stream communities. Freshwater Science 38:1–11.

Metro Vancouver. 2019. EcoHealth Indicators - Canopy Cover and Imperviousness. http://www.metrovancouver.org/data.

Meyer, J. L., M. J. Paul, and W. K. Taulbee. 2005. Stream ecosystem function in urbanizing landscapes. Journal of the North American Benthological Society 24:602–612.

Muotka, T., Paavola, R., Haapala, A., Novikmec, M., and P. Laasonen. 2002. Long-term recovery of stream habitat structure and benthic invertebrate communities from in-stream restoration. Biological Conservation 105:243–253.

Pacific Streamkeepers Federation. 2022, June 6. Pacific Streamkeepers Federation. https://www.pskf.ca/.

Paul, M. J., and J. L. Meyer. 2001. Streams in the Urban Landscape. Annual Review of Ecology and Systematics 32:333–365.

Ranta, E., M. R. Vidal-Abarca, A. R. Calapez, and M. J. Feio. 2021. Urban stream assessment system (UsAs): An integrative tool to assess biodiversity, ecosystem functions and services. Ecological Indicators 121:106980.

R Core Team. 2024. R: A language and environment for statistical computing. R Foundation for Statistical Computing, Vienna, Austria.

Roberge, J.-M., and P. Angelstam. 2004. Usefulness of the umbrella species concept as a conservation tool. Conservation Biology 18:76–85.

Rossi, F., C. Mallet, C. Portelli, F. Donnadieu, F. Bonnemoy, and J. Artigas. 2019. Stimulation or inhibition: Leaf microbial decomposition in streams subjected to complex chemical contamination. Science of the Total Environment 648:1371–1383.

Ruan, X., J. Huang, D. A. R. Williams, K. J. Harker, and S. E. Gergel. 2019. High spatial resolution landscape indicators show promise in explaining water quality in urban streams. Ecological Indicators 103:321–330.

Sudduth, E. B., B. A. Hassett, P. Cada, and E. S. Bernhardt. 2011. Testing the field of dreams hypothesis: functional responses to urbanization and restoration in stream ecosystems. Ecological Applications 21:1972–1988.

Tiegs, S. D., J. E. Clapcott, N. A. Griffiths, and A. J. Boulton. 2013. A standardized cotton-strip assay for measuring organic-matter decomposition in streams. Ecological Indicators 32:131–139.

Walsh, C. J., A. H. Roy, J. W. Feminella, P. D. Cottingham, P. M. Groffman, and R. P. Morgan. 2005. The urban stream syndrome: current knowledge and the search for a cure. Journal of the North American Benthological Society 24:706–723.

Walsh, J. C., K. Connors, E. Hertz, L. Kehoe, T. G. Martin, B. Connors, M. J. Bradford, C. Freshwater, A. Frid, J. Halverson, J. W. Moore, M. H. H. Price, and J. D. Reynolds. 2020a. Prioritizing conservation actions for Pacific salmon in Canada. Journal of Applied Ecology 57:1688–1699.

Walsh, J. C., J. E. Pendray, S. C. Godwin, K. A. Artelle, H. K. Kindsvater, R. D. Field, J. N. Harding, N. R. Swain, and J. D. Reynolds. 2020b. Relationships between Pacific salmon and aquatic and terrestrial ecosystems: implications for ecosystem-based management. Ecology 1010:e03060.

Wang, F., J. Winkler, A. Viña, W. J. McShea, S. Li, T. Connor, Z. Zhao, D. Wang, H. Yang, Y. Tang, J. Zhang, and J. Liu. 2021. The hidden risk of using umbrella species as conservation surrogates: A spatio-temporal approach. Biological Conservation 253:108913.

Warkentin, L., C. Favaro, D. C. Scott, R. Seifert, and J. W. Moore. 2019. Urban planning for fishes: untangling a new project’s effects from old infrastructure and regional patterns. Canadian Journal of Fisheries and Aquatic Sciences 76:378–389.

Webb, J. R., Pearce, N. J. T., Painter, K. J., and A. G. Yates. 2019. Hierarchical variation in cellulose decomposition in least-disturbed reference streams: a multi-season study using the cotton strip assay. Landscape Ecology 34:2353–2369.

Wickham, H., R. François, L. Henry, and K. Müller. 2020. dplyr: A Grammar of Data Manipulation. https://CRAN.R-project.org/package=dplyr

Woodward, G., M. O. Gessner, P. S. Giller, V. Gulis, S. Hladyz, A. Lecerf, B. Malmqvist, B. G. McKie, S. D. Tiegs, H. Cariss, M. Dobson, A. Elosegi, V. Ferreira, M. A. S. Graça, T. Fleituch, J. O. Lacoursière, M. Nistorescu, J. Pozo, G. Risnoveanu, M. Schindler, A. Vadineanu, L. B.-M. Vought, and E. Chauvet. 2012. Continental-scale effects of nutrient pollution on stream ecosystem functioning. Science 336:1438–1440.

Yule, C. M., J. Y. Gan, T. Jinggut, and K. V. Lee. 2015. Urbanization affects food webs and leaf-litter decomposition in a tropical stream in Malaysia. Freshwater Science 34:702– 715.

