## Supplemental Material for "Restoration of urban salmon habitat has limited effects on a key ecosystem function"

### Table of Contents

**SUPPLEMENTARY RESULTS** **p. 1**
Fig. S1 p. 1
Table S1 p. 2
Table S2 p. 5
Table S3 p. 8
**SEM model formulation and selection** p. 17
Fig. S2 p. 20
**SEM diagrams** p. 20

**SUPPLEMENTARY RESULTS** **p. 21**
Fig. S3 p. 21
Fig. S4 p. 22
Fig. S5 p. 23
Fig. S6 p. 24

**LITERATURE CITED** **p. 24**

### SUPPLEMENTARY METHODS

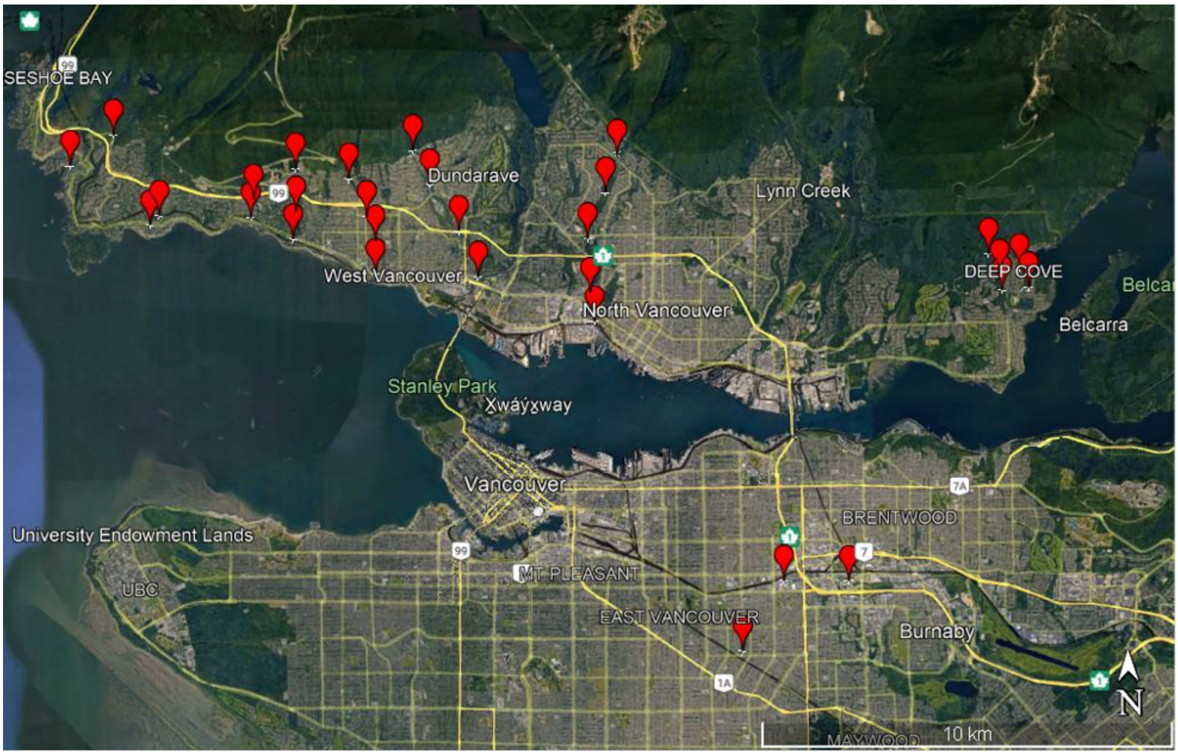


**Figure S1.** Map of study sites (red markers) around the metropolitan Vancouver area.

**Table S1.** Study sites with notes about the variables considered during site selection. Note that PRK-02 was later discarded after concrete blocks were repeatedly washed downstream. For details of calculation of Restoration Effort and Restoration Intensity, refer to Table S3.

| Site ID | Presence of Salmon | Salmon Stocked | Species of Salmon | Impervious Surface (%) | Restoration Effort | Restoration Intensity |
|---|---|---|---|---|---|---|
| BRT01 | YES | NO | Chinook, Chum, Coho, Cutthroat, Pink, Rainbow, Steelhead | 30.82 | 0 | 0 |
| BRT02 | YES | NO | Chinook, Chum, Coho, Cutthroat, Pink, Rainbow, Steelhead | 42.86 | 6 | 13 |
| BRT03 | YES | NO | Chinook, Chum, Coho, Cutthroat, Pink, Rainbow, Steelhead | 28.64 | 0 | 0 |
| BRT04 | YES | NO | Chinook, Chum, Coho, Cutthroat, Pink, Rainbow, Steelhead | 8.06 | 0 | 0 |
| CPS01 | YES | YES | Chum, Coho, Cutthroat, Pink | 34.20 | 1 | 2 |
| CPS02 | YES | YES | Chum, Coho, Cutthroat, Pink | 19.85 | 0 | 0 |
| EAG01 | YES | YES | Chum, Coho, Cutthroat | 32.91 | 4 | 8 |
| EAG04 | YES | YES | Chum, Coho | 3.01 | 0 | 0 |
| GLT01 | YES | NO | Cutthroat | 21.60 | 1 | 6 |
| GLT02 | YES | NO | Cutthroat | 0.93 | 2 | 6 |

| Site ID | Presence of Salmon | Salmon Stocked | Species of Salmon | Impervious Surface (%) | Restoration Effort | Restoration Intensity |
|---|---|---|---|---|---|---|
| GLT03 | YES | NO | Cutthroat | 0.54 | 2 | 4 |
| MCD01 | YES | NO | Chum, Coho, Cutthroat, Steelhead | 50.29 | 11 | 21 |
| MCD02 | YES | NO | Steelhead | 33.57 | 1 | 3 |
| MCD03 | YES | NO | Steelhead | 40.54 | 0 | 0 |
| MCD04 | YES | NO | Steelhead | 16.67 | 0 | 0 |
| MSQ01 | YES | NO | Coho, Chum, Pink | 77.96 | 9 | 16 |
| MSQ02 | YES | NO | Coho, Chum, Pink | 26.81 | 3 | 5 |
| MSQ03 | YES | NO | Coho, Chum, Pink | 34.98 | 5 | 14 |
| MSQ04 | NO | NO | Coho, Chum, Pink | 7.58 | 9 | 12 |
| MSQ05 | NO | NO | Coho, Chum, Pink | 9.79 | 0 | 0 |
| PRK01 | YES | NO | Coho | 10.91 | 4 | 5 |
| PRK02 | YES | NO | Cutthroat | 5.41 | 0 | 0 |
| RGR01 | YES | YS | Chum, Coho, Cutthroat, Pink | 28.82 | 8 | 14 |
| RGR02 | YES | YES | Chum, Coho, Cutthroat, Pink | 18.48 | 1 | 1 |

| Site ID | Presence of Salmon | Salmon Stocked | Species of Salmon | Impervious Surface (%) | Restoration Effort | Restoration Intensity |
|---|---|---|---|---|---|---|
| RGR03 | YES | YES | Chum, Coho, Cutthroat, Pink | 12.13 | 0 | 0 |
| STL02 | YES | NO | Chum | 67.99 | 2 | 5 |
| STL03 | YES | NO | Chum | 91.82 | 7 | 8 |
| STL05 | YES | NO | Chum | 42.79 | 15 | 4 |
| WTM01 | NO | NO | Stocked with Coho | 31.47 | 0 | 0 |
| WTM02 | NO | NO | Stocked with Coho | 30.11 | 0 | 0 |

**Table S2.** Habitat characteristics of study sites.

| Site ID | Riparian Category | Dominant Tree | Dominant Shrub | Dominant Understory | Channel Morphology |
|---|---|---|---|---|---|
| BRT01 | Shrub | Alder | Salmonberry | Ivy | Plane-bed |
| BRT02 | Forest | Alder | Salmonberry | grass | Step-pool |
| BRT03 | Meadow | Cedar | none | buttercups | Cascade |
| BRT04 | Forest | Alder | huckleberry | moss | Cascade |
| CPS01 | Forest | Cedar | Salmonberry | Ivy | Artificial plane-bed |
| CPS02 | Forest | Bigleaf Maple | sword fern | moss | Cascade |
| EAG01 | Forest | Bigleaf Maple | sword fern | moss | Pool-riffle |
| EAG04 | Forest | Bigleaf Maple | sword fern | moss | Pool-riffle |
| GLT01 | Shrub | Alder | Pacific Ninebark | bare | Step-pool |
| GLT02 | Forest | Cedar | sword fern | bare | Cascade |
| GLT03 | Shrub | Western hemlock | Salmonberry | sword fern | Pool-riffle |
| MCD01 | Forest | Alder | Laurel | bare | Artificial plane-bed |
| MCD02 | Forest | Bigleaf Maple | blackberry | sword fern | Cascade |

| Site ID | Riparian Category | Dominant Tree | Dominant Shrub | Dominant Understory | Channel Morphology |
|---|---|---|---|---|---|
| MCD03 | Forest | Cedar | Laurel | Ivy | Cascade |
| MCD04 | Forest | Alder | Salmonberry | moss | Step-pool |
| MSQ01 | Forest | Alder | Salmonberry | Long-stemmed grasses | Pool-riffle |
| MSQ02 | Forest | Alder | Salmonberry | moss | Plane-bed |
| MSQ03 | Forest | Bigleaf Maple | sword fern | Ivy | Cascade |
| MSQ04 | Forest | Alder | Salmonberry | bare | Step-pool |
| MSQ05 | Forest | Vine maple | sword fern | Ivy | Plane-bed |
| PRK01 | Forest | Vine maple | sword fern | Ivy | Plane-bed |
| RGR01 | Forest | Cedar | Laurel | Ivy | Cascade |
| RGR02 | Forest | Cedar | sword fern | Ivy | Step-pool |
| RGR03 | Forest | Cedar | sword fern | bare | Cascade |
| STL02 | Shrub | NA | red elderberry | Canary grass | Dune-ripple |
| STL03 | Shrub | Alder | Pacific Ninebark | bare | Plane-bed |
| STL05 | Forest | Alder | Salmonberry | Ivy | Pool-riffle |

| Site ID | Riparian Category | Dominant Tree | Dominant Shrub | Dominant Understory | Channel Morphology |
|---|---|---|---|---|---|
| WTM01 | Forest | Cedar | bamboo | Ivy | Cascade |
| WTM02 | Forest | Cedar | none | Ivy | Cascade |

**Table S3.** Restoration projects completed in study streams, and points that each project accrued towards the Restoration Effort and Restoration Intensity metrics. For Restoration Effort, projects received 1 point for each project-year of in-stream restoration and 1 point for each project-year of riparian restoration. For Restoration Intensity, projects were assigned as either lightly-assisted, moderately-assisted, or heavily-assisted in line with the suggested classification in Chazdon et al. (2021).

| Stream | Date | Location | Type | Restoration Effort (Points) | Restoration Intensity | Sources | Sites at Location |
|---|---|---|---|---|---|---|---|
| Eagle Creek | 2017 | Near estuary, under driveways | In-stream (shortcrete and plastic baffles added in culvert) | 1 | Moderate | WVSK, 2021 | EAG01 |
| Eagle Creek | 2010 | Marine Drive | In-stream (culvert under road replaced with natural bottom culvert) | 1 | Moderate | WVSK, 2021 | NA |
| Eagle Creek | 2000 | Estuary | In-stream (baffles added in culvert near mouth and rock steps reworked at mouth) | 2 | Intensive | PSKF, 2002; WVSK, 2021 | EAG01 |
| Eagle Creek | 1999 | Eagle Lake | In-stream (diversion pipes installed to augment low summer flow) | 1 | Moderate | PSKF, 2002; WVSK, 2021 | NA |
| Eagle Creek | 1979 | Creek Mouth | In-stream (series of drops and pools created at mouth) | 1 | Intensive | PSKF, 2002; WVSK, 2021 | EAG01 |
| Cypress Creek | 2011 | Intake weir from CAER | In-stream (removeable ladder for late summer to early fall installed) | 1 | Moderate | WVSK, 2021 | CPS01 |

| Stream | Date | Location | Type | Restoration Effort (Points) | Restoration Intensity | Sources | Sites at Location |
|---|---|---|---|---|---|---|---|
| Cypress Creek | 1978 | Unknown | In-stream (incubation box installed for coho salmon) | 1 | Light | PSKF, 2002 | NA |
| Rogers Creek | 2015 | Estuary | In-stream (defined creek path in estuary) | 1 | Intensive | WVSK, 2021 | RGR01 |
| Rogers Creek | 2020 | Estuary | In-stream (estuary reworked to improve pathway through intertidal zone) | 1 | Intensive | WVSK, 2021 | RGR01 |
| Rogers Creek | 1998 | Bellevue Ave | In-stream (baffles constructed under road then upgraded twice) | 3 | Moderate | PSKF, 2002; WVSK, 2021 | RGR01 |
| Rogers Creek | 2003 | Railway Bridge | In-stream (baffles constructed through metal culvert) | 1 | Moderate | PSKF, 2002; WVSK, 2021 | RGR01 |
| Rogers Creek | 2001 | Marine Drive | In-stream (concrete fish ladder installed) | 1 | Moderate | PSKF, 2002; WVSK, 2021 | RGR01 |
| Rogers Creek | Un-known | Marine Drive | In-stream (baffles constructed through culvert) | 1 | Moderate | PSKF, 2002; WVSK, 2021 | RGR01 |
| Rogers Creek | Before 1990 | Palmerston Avenue | In-stream (incubation box installed) | 1 | Light | WVSK, 2021 | RGR02 |
| McDonald Creek | 2014 | Estuary | In-stream (defined creek path in estuary) | 1 | Intensive | WVSK, 2021 | MCD01 |

| Stream | Date | Location | Type | Restoration Effort (Points) | Restoration Intensity | Sources | Sites at Location |
|---|---|---|---|---|---|---|---|
| McDonald Creek | 2001 | Estuary to above Marine Drive | In-stream (baffles installed in open box culvert) | 1 | Moderate | WVSK, 2021 | MCD01 |
| McDonald Creek | Before 1995 | Memorial Park | In-stream (two concrete fish ladders installed) | 2 | Moderate | WVSK, 2021 | MCD01 |
| McDonald Creek | 2012 | Memorial Park | In-stream (rearing pond constructed) | 1 | Intensive | WVSK, 2021 | MCD01 |
| McDonald Creek | 2015 | Fulton Avenue | In-stream (deepened plunge pool, boulders added down-stream to raise water level) | 1 | Intensive | WVSK, 2021 | MCD01 |
| McDonald Creek | Un-known | Fulton Avenue | In-stream (baffles in culvert) | 1 | Moderate | WVSK, 2021 | MCD01 |
| McDonald Creek | Un-known | Fulton Avenue | In-stream (timbers in guides to direct flow away from diversion) | 1 | Moderate | WVSK, 2021 | MCD01 |
| McDonald Creek | 2003 | Hay Park | In-stream (rearing pond constructed) | 1 | Intensive | WVSK, 2021 | MCD02 |
| McDonald Creek | Un-known | Above Highway 1 | In-stream (timbers in guides) | 1 | Moderate | WVSK, 2021 | NA |
| McDonald Creek | 2001 | Marine Drive | In-stream (concrete baffles in culvert) | 1 | Moderate | WVSK, 2021 | MCD01 |

| Stream | Date | Location | Type | Restoration Effort (Points) | Restoration Intensity | Sources | Sites at Location |
|---|---|---|---|---|---|---|---|
| McDonald Creek | 2001 | Estuary to Memorial Park | In-stream (two fish ladders installed in flume and box culverts) | 2 | Moderate | PSKF, 2002; WVSK, 2021 | MCD01 |
| Brothers Creek | Un-known | Taylor Way | In-stream (baffles in culvert) | 1 | Moderate | WVSK, 2021 | BRT02 |
| Brothers Creek | 1997 | Upper Levels Highway | In-stream (baffles in culvert) | 1 | Moderate | WVSK, 2021 | BRT02 |
| Brothers Creek | 2018 | Upper Levels Highway | In-stream (baffles in culvert replaced with concrete version) | 1 | Moderate | WVSK, 2021 | BRT02 |
| Brothers Creek | 1998 | Unknown | In-stream (side channel dug) | 1 | Intensive | PSKF, 2002 | NA |
| Brothers Creek | 1998 | Near Westcott elementary school | In-stream (Westcott pond and fishway constructed) | 1 | Intensive | PSKF, 2002 | BRT02 |
| Brothers Creek | 1997 | Wildwood Lane | In-stream (fishway constructed in culvert) | 1 | Moderate | PSKF, 2002 | BRT02 |
| Brothers Creek | 1977 | Upper Levels Highway 1 | In-stream (timber bolted in herringbone pattern under rds.) | 1 | Moderate | PSKF, 2002 | BRT02 |
| Gallant Creek | Un-known | Indian River Drive | In-stream (box culverts) | 1 | Moderate | PSKF, 2002 | GLT03 |
| Gallant Creek | 2016 | Indian River Drive | In-stream (Repaired culverts) | 1 | Moderate | NSSK, 2019 | GLT03 |

| Stream | Date | Location | Type | Restoration Effort (Points) | Restoration Intensity | Sources | Sites at Location |
|---|---|---|---|---|---|---|---|
| Gallant Creek | 2018 | Cliffwood Road cull-de-sac | In-stream (creek construction to reduce sediment loads) | 1 | Intensive | QM Envir. Services, 2021 | GLT01, GLT02 |
| Gallant Creek | 2015 | Deep Cove Road | Bank stabilization | 1 | Intensive | NSSK, 2019 | GLT01, GLT02 |
| Parkside Creek | 2013 | Cove Cliff Elementary | In-stream (construction of a side channel) | 1 | Intensive | PSKF, 2002 | PRK01 |
| Parkside Creek | 2013 | Cove Cliff Elementary | In-stream (incubation boxes installed) | 1 | Light | PSKF, 2002 | PRK01 |
| Parkside Creek | Un-known | Cove Cliff Elementary | In-stream/Riparian (stream cleanups) | 2 | Light | PSKF, 2002 | PRK01 |
| Parkside Creek | 2015 | Myrtle Park | In-stream (creek un-channelize) | 1 | Intensive | Manley, 2015 | NA |
| Parkside Creek | 2015 | Myrtle Park | Riparian (native riparian vegetation planted) | 1 | Moderate | Manley, 2015 | NA |
| Parkside Creek | 2012 (or prior) | Unknown | Riparian (damaged/exposed section was planted and restored) | 1 | Moderate | NSSK, 2019 | NA |
| Parkside Creek | 2014 | Unknown | Riparian (planting along creek) | 1 | Moderate | NSSK, 2019 | NA |
| Parkside Creek | 2014 | Myrtle Park | Riparian (streamside clean-up) | 1 | Light | NSSK, 2019 | NA |

| Stream | Date | Location | Type | Restoration Effort (Points) | Restoration Intensity | Sources | Sites at Location |
|---|---|---|---|---|---|---|---|
| Still Creek | 2020-2021 | Renfrew Ravine | Riparian (removal of invasives, plant native shrubs and trees) | 2 | Moderate | EYA, 2020 | STL05 |
| Still Creek | 2020-2021 | Canadian Tire | Riparian (removal of invasives, plant native shrubs and trees) | 2 | Moderate | EYA, 2020 | STL03 |
| Still Creek | 2009 | Canadian Tire | In-stream/ Riparian (daylighting, creek widening, habitat naturalization) | 3 | Intensive | Hammond et al. 2018; Rousseau & Verde 2018 | STL03 |
| Still Creek | 2007 | Cornett Road 3400 block | In-stream/ Riparian (reconstructed creek bed, added stream complexity, planted riparian native species) | 2 | Intensive | Rousseau & Verde 2018; Munro & Schwark 2014 | STL03 |
| Still Creek | 2008 | Nootka and Grandview Highway | In-stream/ Riparian (enhancement of habitat and creek bed) | 2 | Intensive | Rousseau & Verde, 2018 | NA |
| Still Creek | 2004-2017 | Renfrew Ravine | Riparian (garbage cleanup, invasive plant removal, native plantings) | 13 | Moderate | Hammond et al., 2018 | STL05 |
| Still Creek | 2005 | Brentwood | Riparian (converting asphalt to wetlands and meadow) | 1 | Intensive | Hammond et al., 2018 | STL02 |

| Stream | Date | Location | Type | Restoration Effort (Points) | Restoration Intensity | Sources | Sites at Location |
|---|---|---|---|---|---|---|---|
| Still Creek | 2005 | Brentwood | Riparian (removing invasive plants, riparian planting) | 1 | Moderate | Hammond et al., 2018 | STL02 |
| Still Creek | 2014 | Burnaby Works Yard | In-stream/ Riparian (restoration in the wetland and creek) | 2 | Intensive | Hammond et al., 2018 | NA |
| Still Creek | 2014 | Burnaby Works Yard | Riparian (invasive plant removal) | 1 | Light | Hammond et al., 2018 | NA |
| Still Creek | 2014 | Burnaby Works Yard | Riparian (installation of bioswales in parking lot) | 1 | Intensive | Hammond et al., 2018 | NA |
| Still Creek | 2006 | Gilmore to Burnaby Lake | Riparian (invasive removal and native planting) | 1 | Moderate | Hammond et al., 2018 | NA |
| Mosquito Creek | 1996-2004 | Marine Drive | In-stream (fish ladder maintenance) | 1 | Moderate | PSKF, 2002 | MSQ01 |
| Mosquito Creek | 1996 | Marine Drive | In-stream (fish ladder installation) | 1 | Moderate | Milne, 2013; PSKF, 2002 | MSQ01 |
| Mosquito Creek | 1998 | William Griffin Park | In-stream (Griffin side channel dug) | 1 | Intensive | Milne, 2013 | MSQ03 |
| Mosquito Creek | 2020 | Bewicke Avenue Bridge | In-stream (adding large log/boulder clusters to remove erosive forces) | 1 | Moderate | NSSK, 2020 | MSQ01 |

| Stream | Date | Location | Type | Restoration Effort (Points) | Restoration Intensity | Sources | Sites at Location |
|---|---|---|---|---|---|---|---|
| Mosquito Creek | 2014 | Bewicke Bridge | In-stream/ Riparian (rehabilitate the shorefront with a series of habitat benches; upgrades to riparian habitat & plantings) | 2 | Intensive | NSSK, 2019 | MSQ01 |
| Mosquito Creek | 1999 | Mosquito Creek Park | In-stream (constructed side channel called Fell Channel) | 1 | Intensive | Milne, 2013 | MSQ04 |
| Mosquito Creek | 1996 | East Bank back channel (possibly near Highway 1) | In-stream (lengthened and deepened back channel to create habitat pool) | 1 | Intensive | Milne, 2013; PSKF, 2002 | NA |
| Mosquito Creek | 1997 | East Bank back channel (possibly near Highway 1) | In-stream/ Riparian (lengthened back channel and planted native plants) | 2 | Intensive | Milne, 2013 | NA |
| Mosquito Creek | 1997 | West bank channel (possibly near Low Level Bridge) | In-stream/ Riparian (back channel lengthened and deepened, native plants planted) | 2 | Intensive | Milne, 2013 | NA |
| Mosquito Creek | 1997, 2003 | Marine drive up to Larson Road | Riparian (native riparian vegetation planted) | 2 | Moderate | Milne, 2004 | MSQ01, MSQ02 |
| Mosquito Creek | 2000, 2001, 2002 | Mosquito Creek Park | Riparian (native riparian vegetation planted) | 3 | Moderate | Milne, 2004; PSKF, 2002 | MSQ04 |

| Stream | Date | Location | Type | Restoration Effort (Points) | Restoration Intensity | Sources | Sites at Location |
|---|---|---|---|---|---|---|---|
| Mosquito Creek | 1995 | East bank (19th street and Bewicke Avenue) | Riparian (slope stabilization and planting) | 1 | Intensive | Milne, 2004; PSKF, 2002 | MSQ02 |
| Mosquito Creek | 2012 or prior | Just above Evergreen Place. | Riparian (tree planting) | 1 | Moderate | NSSK, 2019 | MSQ04 |
| Mosquito Creek | 2013 | Upper Mosquito Creek | Riparian (Installation of anchors and Debris Fence net) | 1 | Moderate | NSSK, 2019 | NA |
| Mosquito Creek | 2012-2015 | Mosquito Creek Park | Riparian (invasive plant removal and native planting) | 3 | Moderate | NSSK, 2019 | MSQ04 |
| Mosquito Creek | 2015 | William Griffin Park | In-stream (Gravel removal from debris basin behind Evergreen Place, placed back in Mosquito Creek at the park to improve in-stream habitat) | 1 | Intensive | NSSK, 2019 | MSQ03 |
| Mosquito Creek | 2014 | William Griffin Park | Riparian (new bioswale/wetland area constructed outside parking lot to filter runoff) | 1 | Intensive | NSSK, 2019 | MSQ03 |
| Mosquito Creek | 2014, 2017 | Fell Intake | In-stream (cleaning of blockages) | 2 | Moderate | NSSK, 2019 | NA |
| Mosquito Creek | 2014 | William Griffin Park | In-stream (improvements to Griffin channel) | 1 | Intensive | NSSK, 2019 | MSQ03 |

| Stream | Date | Location | Type | Restoration Effort (Points) | Restoration Intensity | Sources | Sites at Location |
|---|---|---|---|---|---|---|---|
| Mosquito Creek | 2015 | Mosquito Creek Park | In-stream (weirs built by Fell intake and fell intake itself rebuilt) | 1 | Intensive | NSSK, 2019 | MSQ04 |
| Mosquito Creek | 2016 | Marine Drive Bridge | In-stream (Gravel removal) | 1 | Intensive | NSSK, 2019 | MSQ01 |
| Mosquito Creek | 2017 | 3rd Street Bridge | Riparian (invasive removal and native plant planting) | 1 | Moderate | NSSK, 2019 | MSQ01 |
| Mosquito Creek | 2017, 2018 | William Griffin Park | In-stream (add woody debris to the Griffen Channel) | 1 | Moderate | NSSK, 2019 | MSQ03 |
| Mosquito Creek | 2017 | Unknown | Riparian (cedar tree planting) | 1 | Moderate | Casidy, 2017 | NA |

### SEM Model Formulation and Selection

We chose the piecewise structural equation model (SEM) approach because it is flexible in terms of the structure of the component models, accommodating generalized and mixed-effects models (Lefcheck 2016). The initial SEM model developed based on hypothesized relationships among variables consisted of four linear mixed-effects models:

(1) Impervious surface ~ Elevation + (1 | Watershed)

(2) RI at site ~ Elevation + Impervious surface + (1 | Watershed)

(3) PCA component 1 ~ Elevation + RI at site + (1 | Watershed)

(4) TLDD ~ Elevation + Impervious surface + RI at site + PCA component 1 + riffle + (1 | Site)

Using the ‘DHARMa’ package (Hartig 2022), we found that models 1-3 did not meet the assumption that the correct distribution was used (KS test; model 1 $p = 0.04$, model 2 $p = 0.04$, model 3 $p < 0.001$), and all four models showed significant deviations between the expected quantiles of model predictions and residuals.

Exploring potential reasons for these deviations, we first considered associations between elevation and other variables, which were the reason that we had chosen an SEM approach in the first place. However, our dataset had more data collected from low elevations than high elevations, in part because some streams simply did not originate at particularly high elevation, and in part because access to other streams was challenging close to their headwaters. Furthermore, most variables showed greater variance among low-elevation sites than among high-elevation sites, or in other words, there was a broader range of possible conditions in low-elevation sites than in higher-elevation sites. This is illustrated by the relationship between impervious surface and elevation, which shows heterogeneity of variance with elevation and is possibly nonlinear (Figure S2).

To address this, we switched our component model formulas to models with a dispersion parameter accounting variance with elevation using the glmmTMB package version 1.1.10 (Brooks et al. 2017). In addition, our RI and RE metrics represent counts of restoration projects. Their overdispersion could be addressed by fitting a generalized linear model with the negative binomial distribution. We also tried different random effects structures (e.g. watershed, site, and site nested within watershed) and found that in several cases, assumptions were better met by models without random effects than those with random effects. Fitting different formulations of models using glmmTMB raised p-values for the KS test above the level of $\alpha = 0.05$ for models 1, 2, and 3, and moderately improved the deviation between the expected quantiles of model predictions and residuals in model 4.

Incorporating these glmmTMB models into the piecewise SEM proved challenging, however, for several reasons. The piecewiseSEM package does not support generalized models with the negative binomial family. It also ignores the specification of the dispersion model when getting variances. Thus, for many paths, the standardized coefficients could not be computed.

As a result, we returned to our original model formulation, but are cautious in our interpretation as we recognize that not all model assumptions are met, especially for model 3.

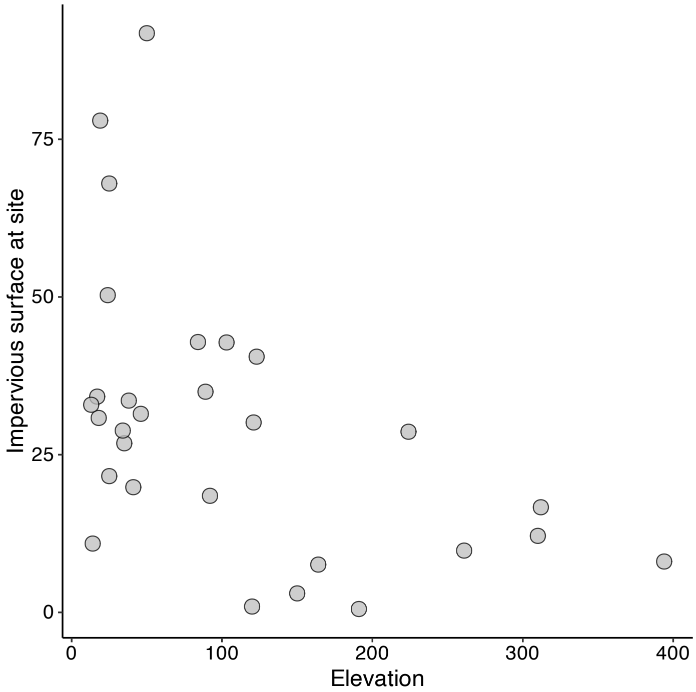


**Fig. S2.** Impervious surface (% of surrounding 400 $m^2$ total area) at study sites of different elevations.

### SEM Diagrams

Plotting in the partialSEM package does not allow the user to change the layout or geometry of SEM diagrams. Therefore, to construct path diagrams that were easier to interpret, we re-ran SEM models using the 'lavaan' package version 0.6-19 (Rosseel 2012), produced diagrams with customized layouts using the 'tidySEM' package version 0.2.7 (van Lissa 2024), and then manually edited path coefficients and path color and symbology based on the output from the piecewiseSEM models.

### SUPPLEMENTARY RESULTS

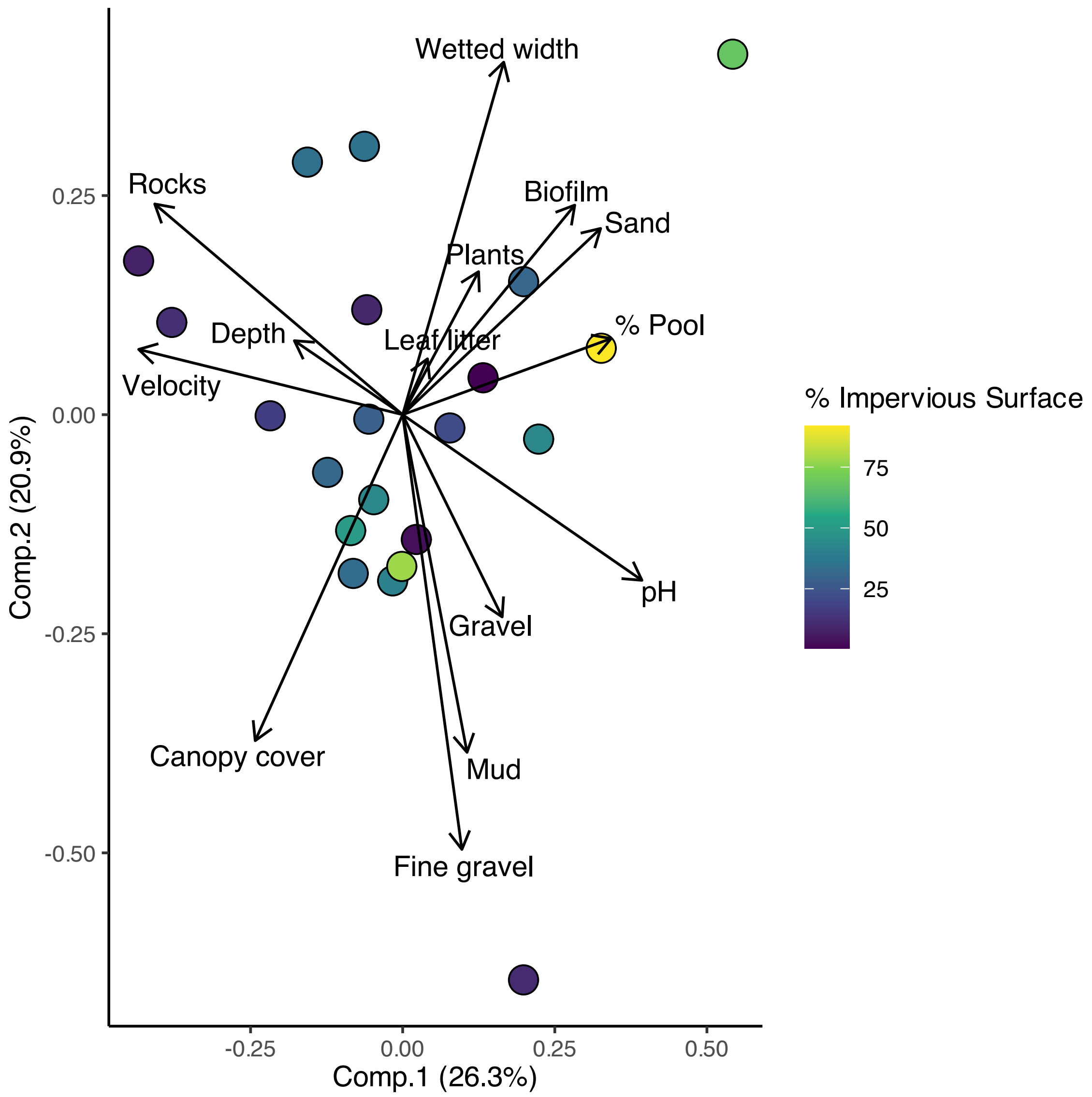


**Fig. S3.** Principal component analysis of habitat variables measured at 21 study sites, including pH as a measure of water chemistry.

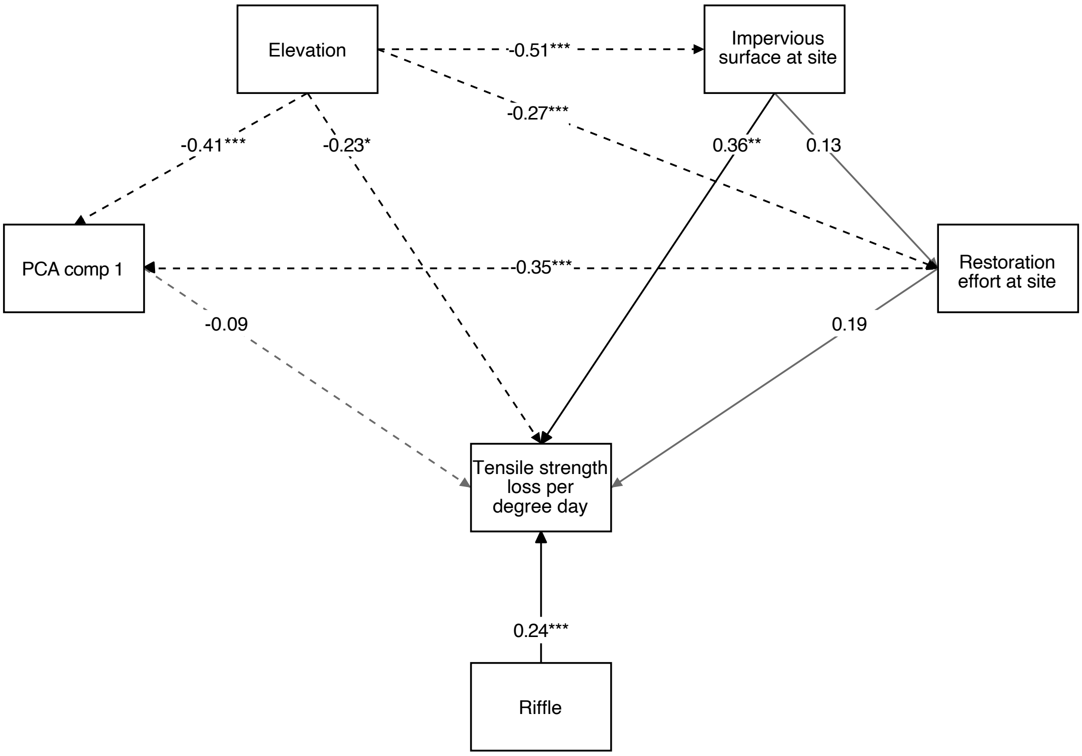


**Fig. S4.** Results of a structural equation model examining the direct and indirect effects of elevation, urbanization (as measured by amount of impervious surface surrounding a stream reach), restoration effort at each stream reach, and habitat characteristics on cotton strip decomposition rates across the early- and mid-summer study periods. Significant ($p < 0.05$) paths are shown with black lines, and non-significant paths ($p > 0.05$) are shown with grey lines. Negative paths are indicated by dashed lines.

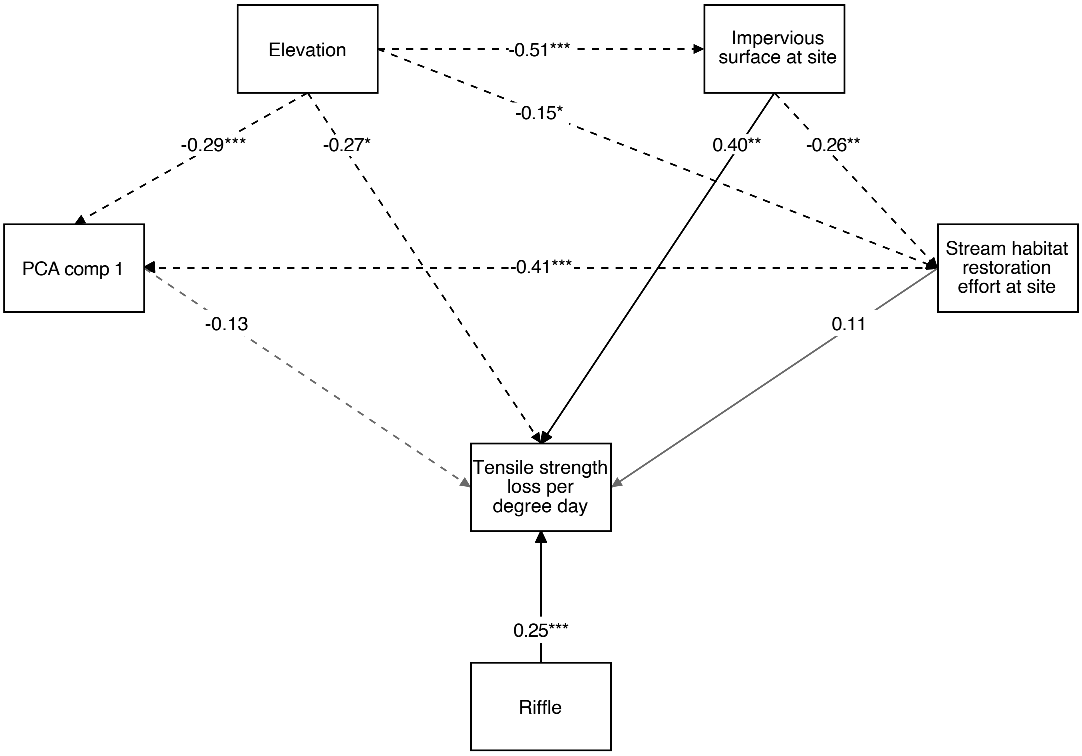


**Fig. S5.** We also calculated a restoration effort metric which only included projects that directly restored stream habitat, excluding projects that focused on fish passage (such as fish ladders) and the construction of spawning habitat outside of the main stream channel (such as ponds). This figure shows results of a structural equation model with the modified restoration effort metric. The model examines the direct and indirect effects of elevation, urbanization (as measured by amount of impervious surface surrounding a stream reach), restoration effort at each stream reach, and habitat characteristics on cotton strip decomposition rates across the early- and mid-summer study periods. Significant ($p < 0.05$) paths are shown with black lines, and non-significant paths ($p > 0.05$) are shown with grey lines. Negative paths are indicated by dashed lines.

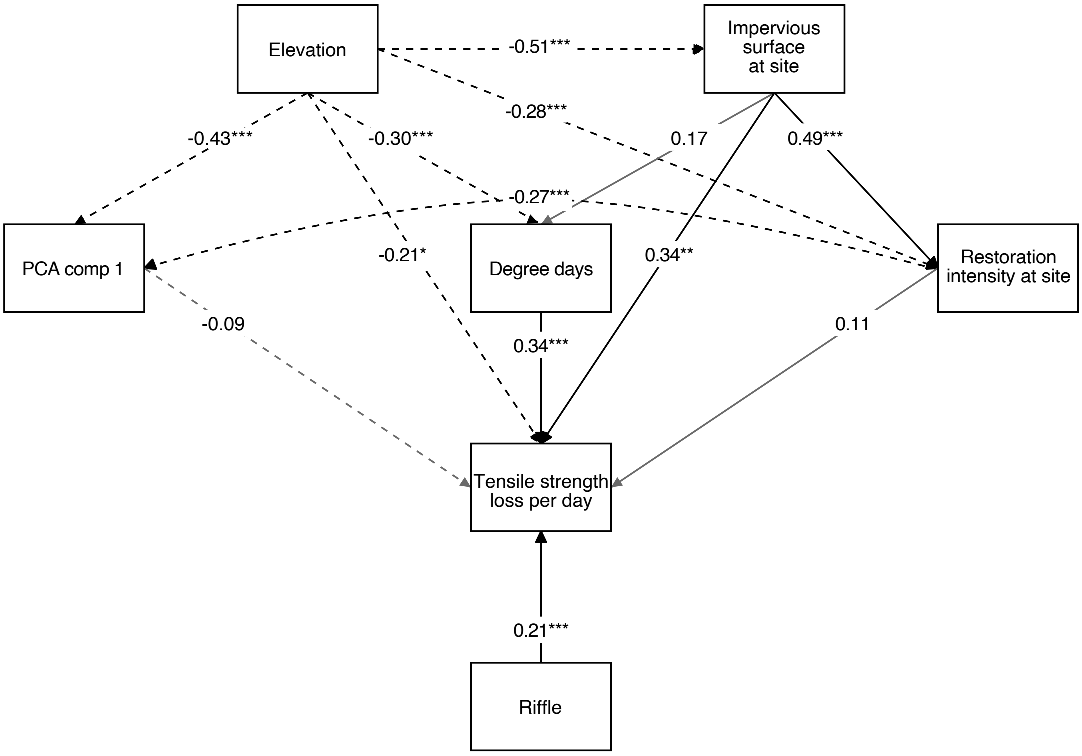


**Fig. S6.** We also constructed a structural equation model using tensile strength loss per day, rather than temperature-adjusted tensile strength loss as in Figure 3. Instead, here we include the number of degree days of warming experienced at the site as an explicit variable in the structural equation model. Thus, this model examines the direct and indirect effects of elevation, urbanization (as measured by amount of impervious surface surrounding a stream reach), temperature, intensity of restoration interventions at each stream reach, and habitat characteristics on cotton strip decomposition rates across the early- and mid-summer study periods. Significant ($p < 0.05$) paths are shown with black lines, and non-significant paths ($p > 0.05$) are shown with grey lines. Negative paths are indicated by dashed lines.

### LITERATURE CITED

Brooks, M., Kristensen, K., van Benthem, K.J., Magnusson, A., Berg, C.W., Nielsen, A., Skaug, H.J., Mächler, M., and B. Bolker. 2017. {glmmTMB} Balances Speed and Flexibility Among Packages for Zero-inflated Generalized Linear Mixed Modeling. The R Journal 9:378–400.

Casidy, K. (2017) *Cedar Tree Planting & Updates.* Mosquito Creek. https://mosquitocreeknv.weebly.com/blog/archives/09-2017. Accessed 8 July 2022.

Chazdon, R.L., Falk, D.A., Banin, L.F., Wagner, M., Wilson, S.J., Grabowski, R.C., & Suding, K.N. (2021) The intervention continuum in restoration ecology: rethinking the active-passive dichotomy. *Restoration Ecology* e13535. DOI: 10.1111/rec.13535.

Environmental Youth Alliance (EYA). (2020). *Still Creek Restoration*. Environmental Youth Alliance. https://eya.ca/project-view/still-creek-restoration/. Accessed 8 July 2022.

Hammond, H., Boyle, C., & Doyle-Yamaguchi, E. (2018). *Still Creek watershed ecosystem-based stewardship plan for ecological restoration.* Silva Forest Foundation.

Lefcheck, J.S. 2016. piecewiseSEM: Piecewise structural equation modeling in R for ecology, evolution, and systematics. Methods in Ecology and Evolution 7:573–579.

Manley, C. (2015). *Parkside creek enhancement & garden 2015 - Deep Cove, BC.* Chad Manley practice in landscape and building arts. https://www.chadmanley.ca/Parkside-Creek-Enhancement-Garden. Accessed 8 July 2022.

Milne, K.R. (2004) *Revegetation - Mosquito Creek*. Mosquito Creek Watershed. https://web.archive.org/web/20041128230308fw_/http://www.mosquitocreek.org/rest

ore_reveg_Mos.html. Accessed 8 July 2022.

Milne, K.R. (2013). *Fish Habitat – Mosquito Creek.* Mosquito Creek Watershed. https://web.archive.org/web/20061126090746/http://www.mosquitocreek.org/restore_fishhab_Mos.html. Accessed 8 July 2022.

Munro, K., & Schwark, k. (2014). *Still Creek enhancement – project funding administrative report.* City of Vancouver. https://council.vancouver.ca/20140916/documents/a9.pdf. Accessed 8 July 2022.

North Shore Streamkeepers (NSSK). (2019) *What We Do*. North Shore Streamkeepers. https://web.archive.org/web/20190914195916/http://www.nssk.ca/what-we-do/; click the dropdown “Archives” tab on the right side and information is in meeting minutes from individual months. Accessed 8 July 2022.

North Shore Streamkeepers (NSSK). (2020). *Lower Mosquito Creek salmon enhancement project 2020.* North Shore Streamkeepers. http://www.nssk.ca/lower-mosquito-creek-salmon-enhancement-project-2020/. Accessed 8 July 2022.

Pacific Streamkeepers Federation (PSKF). (2002) *Watershed profiles*. The Pacific Streamkeepers Federation. http://www.pskf.ca/ecology/watershed/index.html. Accessed 8 July 2022.

QM Environmental Services. (2021). *Gallant creek sediment basin.* QM Environmental Services. https://www.qmenv.com/portfolio-items/gallant-creek-sediment-basin/. Accessed 8 July 2022.

Rousseau, & Verde. (2018). *Still Creek enhancement project – 2900 Nootka Street administrative report* (A12). City of Vancouver. https://vancouver.ca/files/cov/still-creek-2900-nootka-street.pdf. Accessed 8 July 2022.

Rosseel, Y. 2012. lavaan: An R Package for Structural Equation Modeling. Journal of Statistical Software 48:1–36.

van Lissa, C.J. 2024. tidySEM: Tidy Structural Equation Modeling. https://CRAN.R-project.org/package=tidySEM

West Vancouver Streamkeepers (WVSK). (2021). *Creek Data*. The West Vancouver Streamkeepers Society. https://www.westvancouverstreamkeepers.ca/creek-data. Accessed 8 July 2022.
